# Evolutionary history of the porpoises (*Phocoenidae*) across the speciation continuum: a mitogenome phylogeographic perspective

**DOI:** 10.1101/851469

**Authors:** Yacine Ben Chehida, Julie Thumloup, Cassie Schumacher, Timothy Harkins, Alex Aguilar, Asunción Borrell, Marisa Ferreira, Lorenzo Rojas-Bracho, Kelly M. Roberston, Barbara L. Taylor, Gísli A. Víkingsson, Arthur Weyna, Jonathan Romiguier, Phillip A. Morin, Michael C. Fontaine

## Abstract

Historical changes affecting food resources are a major driver of cetacean evolution. Small cetaceans like porpoises (Phocoenidae) are among the most metabolically challenged marine mammals and are particularly sensitive to changes in their food resources. The seven species of this family inhabit mostly temperate waters and constitute a textbook example of antitropical distribution. Yet, their evolutionary history remains poorly known despite major conservation issues threatening the survival of some porpoises (e.g., vaquita and Yangzte finless porpoises). Here, we reconstructed their evolutionary history across the speciation continuum, from intraspecific subdivisions to species divergence. Phylogenetic analyses of 63 mitochondrial genomes suggest that, like other toothed whales, porpoises radiated during the Pliocene in response to deep environmental changes. However, all intra-specific phylogeographic patterns were shaped during the Quaternary Glaciations. We observed analogous evolutionary patterns in both hemispheres associated with convergent adaptations to coastal *versus* oceanic environments. This result suggests that the mechanism(s) driving species diversification in the relatively well-known species from the northern hemisphere may apply also to the poorly-known southern species. In contrast to previous studies, we showed that the spectacled and Burmeister’s porpoises share a more recent common ancestor than with the vaquita that diverged from southern species during the Pliocene. The low genetic diversity observed in the vaquita carried signatures of a very low population size throughout at least the last 5,000 years, leaving one single relict mitochondrial lineage. Finally, we observed unreported subspecies level divergence within Dall’s, spectacled and Pacific harbor porpoises, suggesting a richer evolutionary history than previously suspected. These results provide a new perspective on the mechanism driving the adaptation and speciation processes involved in the diversification of cetacean species. This knowledge can illuminate their demographic trends and provide an evolutionary framework for their conservation.

## 1. Introduction

Most cetaceans possess a tremendous potential for dispersal in an environment that is relatively unobstructed by geographical barriers. This observation begs the question of how do populations of such highly mobile pelagic species in such an open environment split and become reproductively isolated from each other to evolve into new species. Recent micro- and macro-evolutionary studies showed that changes in environmental conditions (Steeman et al., 2009), development of matrilinearly transmitted cultures (Whitehead, 1998), and resource specialization (Fontaine et al., 2014; Foote et al., 2016; Louis et al., 2014) are major drivers of population differentiation and speciation in cetacean species. Yet, the processes that link these two evolutionary timescales are still not fully understood and empirical examples are limited (Foote et al., 2016; Steeman et al., 2009).

Several cetacean taxa display an antitropical distribution where the distribution of closely related taxa occurs on either side of the equator but are absent from the tropics (Barnes, 1985; Hare et al., 2002; Pastene et al., 2007). Multiple mechanisms have been proposed to explain such a peculiar distribution (Burridge, 2002). In cetaceans, the predominant hypothesis implies dispersal and vicariance of temperate species enabled by climatic fluctuations, especially during the Quaternary glacial-interglacial cycles (Davies, 1963). During glacial periods, cold adapted taxa extended their range into the tropics and possibly crossed the equator. In the subsequent warmer interglacial periods, these taxa would shift their range to higher latitudes. This geographic isolation in both hemispheres promoted allopatric divergence of conspecific taxa, resulting in their antitropical distribution. A closely related scenario suggests that the rise of the overall sea temperature during interglacial periods would have altered the winds direction and upwelling strength, leading to a redistribution of feeding areas for cetaceans toward higher latitudes, which in turn promoted their antitropical distribution (Pastene et al., 2007). Another plausible hypothesis implies that broadly distributed species, such as several cetacean species, were outperformed in the tropics by more competitive species (Burridge, 2002). A combination of these different mechanism is also possible.

The porpoises family (Phocoenidae) displays one of the best known example of antitropical distribution (Barnes, 1985). Porpoises are among the smallest cetaceans and represent an interesting evolutionary lineage within the Delphinoidea, splitting from the Monodontidae during the Miocene (∼15 Million years ago) (McGowen et al., 2009; Steeman et al., 2009). Gaskin (1982) suggested that porpoises originated from a tropical environment and then radiated into temperate zones in both hemispheres. In contrast, based on the location of the oldest fossils, Barnes (1985) suggested that they arose in a more temperate environment of the North Pacific Ocean and subsequently colonized the southern hemisphere, the Indian and Atlantic Oceans. Porpoises consist of seven extant species that occur in both hemispheres in pelagic, coastal, and riverine habitats (Fig. 1a). The family includes primarily cold-water tolerant species, but two sister species of finless porpoises (*Neophocoena phocaenoides, N. asiaeorientalis*) (Zhou et al., 2018) inhabit the tropical waters of the Indo-Pacific preferring mangrove zones. They are also found in the Yellow Sea, East China Sea, Sea of Japan and in estuaries and large river systems of the Indus, Ganges, and Yangtze rivers. The remaining species are considered cold-water tolerant and are antitropically distributed. In the Northern Hemisphere, the harbor porpoise (*Phocoena phocoena*) inhabits the coastal waters of the North Pacific and North Atlantic, while the Dall’s porpoise (*Phocoenoides dalli*) occupies the oceanic waters of the North Pacific. This neritic-oceanic habitat segregation is mirrored in the southern hemisphere with the Burmeister’s porpoise (*Phocoena spinipinnis*), occupying the coastal waters of South America and the poorly known spectacled porpoise (*Phocoena dioptrica*) occupying the circum-Antarctic oceanic waters. The vaquita depart from the other species with an extremely localized geographical distribution in the upper Gulf of California and is now critically endangered (Morell, 2017).

**Fig. 1.**
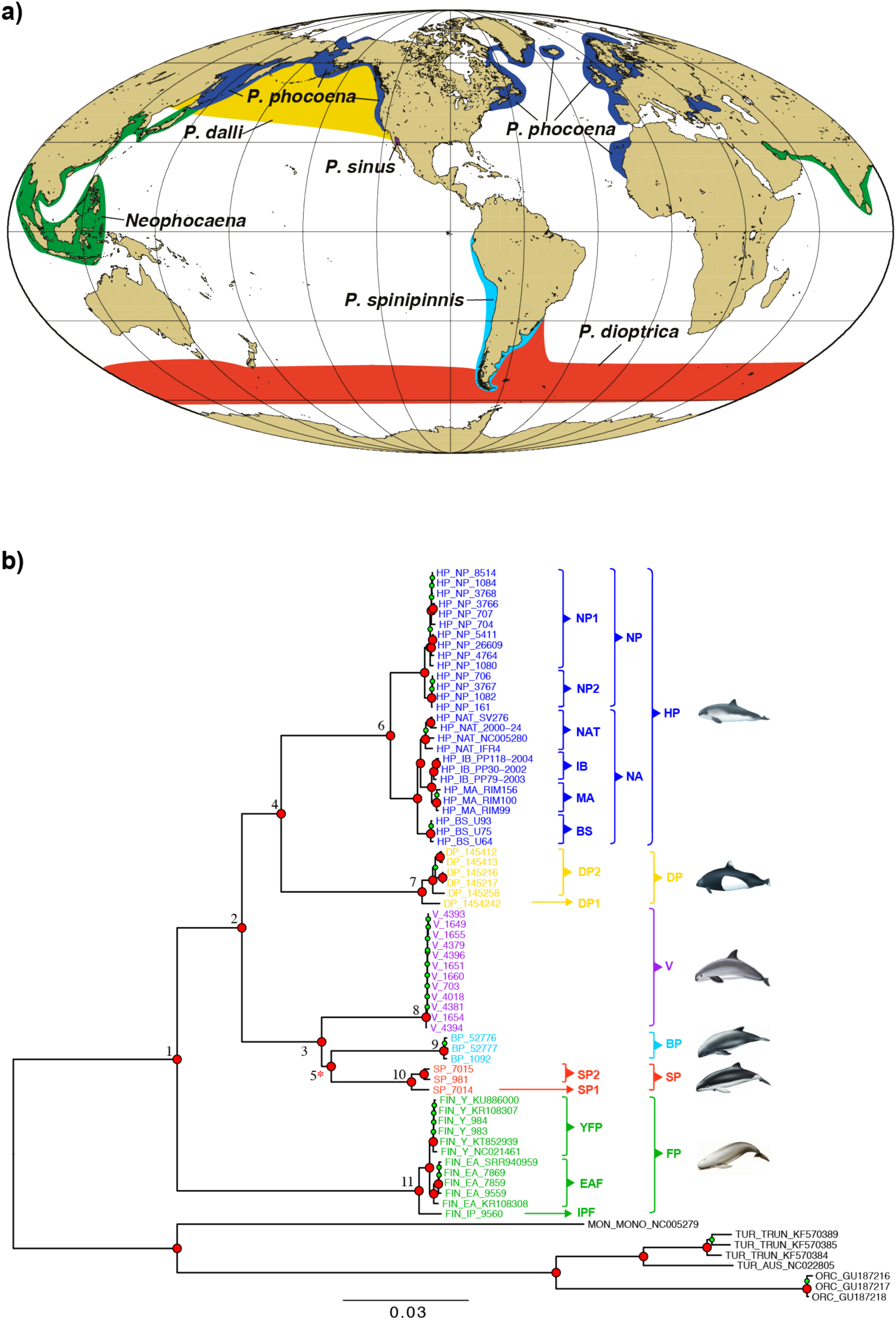
(a) Geographical range of the seven porpoise species modified from Berta et al. (2015). (b) Maximum-likelihood mitochondrial phylogeny. The external branches and the tip labels are colored by species. The tree is rooted with eight sequences from four closely related *Odontoceti* species (in black). Numbers at the nodes are discussed in the text. The nodes indicated in red and green represent nodes with bootstrap support ≥ 90% and ≥70%, respectively. The red star indicates a node with a 63% bootstrap support in the Neighbor-joining tree (Fig. S2c). The acronyms are provided in Table 1. The scale bar unit is in substitution per site.

**Table 1.** Taxon sample size (n) and descriptive statistics for the shotgun sequencing and mitochondrial assembly per species and mitochondrial lineage. The statistics include the total number of reads before and after filtering (R*_b_* and R*_a_*), the sequencing coverage depth, the size of the mitochondrial assembly (in base-pairs), and the GC content in percent (%GC). The mean value and the standard deviation are shown.

| Lineage | n | $R_b$ | $R_a$ | Depth | Assembly size | % GC |
| --- | --- | --- | --- | --- | --- | --- |
| Harbor porpoise (HP) | 27 | 30.8 ± 28.3 | 28.5 ± 25.9 | 1323.9 ± 1800.0 | 16383.8 ± 0.8 | 40.6 ± 0.1 |
| NP1 | 10 | 5.1 ± 1.8 | 5.0 ± 1.7 | 99.6 ± 92.6 | 16384.3 ± 0.5 | 40.7 ± 0.0 |
| NP2 | 4 | 5.9 ± 1.3 | 5.6 ± 1.2 | 97.2 ± 119.8 | 16384.0 ± 0.0 | 40.7 ± 0.0 |
| NAT | 4 | 56.9 ± 3.4 | 52.2 ± 2.9 | 1656.3 ± 1058.9 | 16383.3 ± 0.6 | 40.6 ± 0.0 |
| MA | 3 | 59.8 ± 5.4 | 54.9 ± 5.2 | 1431.0 ± 648.1 | 16384.0 ± 0.0 | 40.6 ± 0.0 |
| IB | 3 | 65.2 ± 8.2 | 60.5 ± 7.6 | 3169.7 ± 1882.6 | 16384.0 ± 0.0 | 40.5 ± 0.0 |
| BS | 3 | 60.0 ± 2.2 | 54.8 ± 1.3 | 4755.3 ± 1376.9 | 16382.0 ± 0.0 | 40.6 ± 0.0 |
| Dall's porpoise (DP) | 6 | 2.8 ± 0.9 | 2.7 ± 0.9 | 49.5 ± 14.5 | 16367.5 ± 0.8 | 40.5 ± 0.1 |
| DP1 | 1 | 4.5 | 4.4 | 74.0 | 16369.0 | 40.6 |
| DP2 | 5 | 2.5 ± 0.5 | 2.4 ± 0.5 | 44.6 ± 9 | 16367.2 ± 0.4 | 40.5 ± 0.0 |
| Vaquita (V) | 12 | 5.8 ± 5.7 | 5.7 ± 5.5 | 113.3 ± 108.6 | 16370.0 ± 0.0 | 39.8 ± 0.0 |
| Burmeister's porpoise (BP) | 3 | 2.3 ± 0.3 | 2.3 ± 0.3 | 125.0 ± 121.2 | 16378.7 ± 8.1 | 40.1 ± 0.0 |
| Spectacled porpoise (SP) | 3 | 2.4 ± 0.2 | 2.3 ± 0.2 | 55.0 ± 30.5 | 16371.0 ± 0.0 | 39.7 ± 0.1 |
| SP1 | 1 | 2.2 | 2.1 | 41.0 | 16371.0 | 39.6 |
| SP2 | 2 | 2.5 ± 0.0 | 2.4 ± 0.0 | 62.0 ± 39.6 | 16371.0 ± 0.0 | 39.7 ± 0.1 |
| Finless porpoises (FP) | 12 | 2.7 ± 1.1 | 2.6 ± 1.1 | 92.9 ± 74.9 | 16383.1 ± 5.8 | 40.8 ± 0.0 |
| IPF | 1 | 2.3 | 2.2 | 77.0 | 16385.0 | 40.8 |
| EAF | 5 | 3.0 ± 1.5 | 2.9 ± 1.4 | 53.2 ± 22.8 | 16381.2 ± 7.5 | 40.0 ± 0.0 |
| YF | 6 | 2.3 ± 0.3 | 2.2 ± 0.3 | 180.0 ± 101.8 | 16386.0 ± 0.0 | 40.8 ± 0.0 |

The lineage code includes the vaquita (*P. sinus*, V), Burmeister's porpoise (*P. spinipinnis*, BP), spectacled porpoise (*P. dioptrica*, SP), Dall's porpoise (*P. dalli*, DP), harbor porpoise (*P. phocoena*, HP) and finless porpoise (*N. phocaenoides* + *N. asiaeorientalis*, FP). Some species have been also further subdivided into distinct groups: harbor porpoises are divided into North Pacific (NP) and North Atlantic (NA), and within each of these groups, further subdivisions are recognized. Four groups are recognized within NA: NAT (North Atlantic), IB (Iberian), MA (Mauritanian), BS (Black Sea). NP is divided into two subgroups: NP1 and NP2. Three sub-species or species are also recognized among the finless porpoises: Indo-Pacific (IPF), Yangtze finless (YFP) and East Asia finless (EAF). Spectacled porpoise subgroups are designated as SP1 and SP2. Dall's porpoise subgroups as DP1 and DP2.

With the exception of vaquitas, all species of porpoises exhibit a relatively broad distribution range that appear fairly continuous at a global scale. Nevertheless, despite having the ability to disperse over vast distance in an open environment, many include distinct subspecies, ecotypes, or morphs. For example, the finless porpoises not only include two recognized species, but also an incipient species within the Yangtze River (Chen et al., 2017; Zhou et al., 2018); the harbor porpoise also displays a disjunct distribution with at least four sub-species reported (Fontaine, 2016); at least two forms of Dall’s porpoise have been described (Hayano et al., 2003); and the Burmeister’s porpoise also shows evidence of deep population structure (Rosa et al., 2005). Such intraspecific subdivisions, also observed in killer whales (Foote et al., 2016) and dolphins (Louis et al., 2014), illustrate the evolutionary processes in action, and can, in some cases, eventually lead to new species. Porpoises are thus an interesting model to investigate the evolutionary processes at both micro- and macroevolutionary time scale to better understand present and historical mechanisms governing population structure, adaptation to different niches, and speciation.

From a conservation perspective, the coastal habitat of many porpoise species largely overlaps with areas where human activities are the most intense. These can have dramatic impacts on natural populations. For example, the vaquita lost 90% of its population between 2011 and 2016 leaving about 30 individuals in 2017 (Thomas et al., 2017), and less than 19 in 2019 (Jaramillo Legorreta et al., 2019). This species is on the brink of extinction and currently represents the most endangered marine mammal. Finless porpoise also faces major conservation issues, especially the lineage within the Yangtze River (*N. a. asiaeorientalis*) in China, also on the brink of extinction due to human activities (Wang et al., 2013; Wang and Reeves, 2017). Similarly, several populations of harbor porpoises are highly threatened (Birkun and Frantzis, 2008; Read et al., 2012). Little information about the spectacled and Burmeister’s porpoises is available. While anthropogenic activities are an undeniable driver of the current threats to biodiversity, the evolutionary context should also be considered when assessing their vulnerability (Dufresnes et al., 2013). Such knowledge is useful to characterize population dynamics, identify evolutionary significant units relevant for conservation, recent or historical split related to environmental variation, evolutionary or demographic trends, and evolutionary processes that could explain or enhance or mitigate the current threats experienced by species (Malaney and Cook, 2013; Moritz and Potter, 2013).

Porpoise evolutionary history and biogeography remains contentious and superficial (Fajardo-Mellor et al., 2006). Previous phylogenetic studies led to incongruent results, as there are disagreements regarding some of their relationships, in particular about the position of the vaquita, Dall’s, Burmeister’s and spectacled porpoises (Barnes, 1985; Rosel et al., 1995b; Fajardo-Mellor et al., 2006; McGowen et al., 2009). So far, molecular phylogenetic relationships among porpoises have been estimated using short sequences of the D-loop and cytochrome *b* (McGowen et al., 2009; Rosel et al., 1995b). However, the D-loop can be impacted by high levels of homoplasy that blurs the resolution of the tree (Torroni et al., 2006) and the cytochrome *b* may have limited power to differentiate closely related taxa (Viricel and Rosel, 2011).

In this study, we sequenced and assembled the whole mitogenome from all extant porpoise species, including most of the known lineages within species to resolve their phylogenetic relationships and reconstruct their evolutionary history. More specifically, (1) we assessed the phylogenetic and phylogeographic history of the porpoise family based on the whole mitogenomes including the timing and tempo of evolution among lineages; (2) we assessed the genetic diversity among species and lineages and (3) reconstructed the demographic history for some lineages for which the sample size was suitable. (4) We placed the evolutionary profile drawn for each lineage and species into the framework of past environmental changes to extend our understanding of porpoise biogeography. Finally, (5) we interpreted the IUCN conservation status of each taxa in the light of their evolutionary history.

## 2. Material & methods

### 2.1. Data collection

Porpoise tissue samples from 56 live-biopsy or dead stranded animals (Table S1) were stored in salt-saturated 20% DMSO or 95% Ethanol and stored at −20°C until analyses. Genomic DNA was extracted from tissues using the *PureGene* or *DNeasy* Tissue kits (Qiagen), following the manufacturer’s recommendations. DNA quality and quantity were assessed on an agarose gel stained with ethidium bromide, as well as using a Qubit-v3 fluorometer. Genomic libraries for 44 porpoise samples including 3 spectacled porpoises, 3 Burmeister’s porpoises, 12 vaquita, 6 Dall’s porpoises, 3 East Asian finless porpoises, 2 Yangtze finless, 1 Indo-Pacific finless and 14 North Pacific harbor porpoises. Libraries were prepared by Swift Biosciences Inc. using either the Swift Biosciences Accel-NGS double-strand 2S (harbor porpoises) or single-strand 1S Plus DNA Library Kit (all other species), following the user protocol and targeting an insert-size of ∼350 base-pairs (bps). The libraries were indexed and pooled in groups of 2 to 12 for paired-end 100 bps sequencing in five batches on an Illumina MiSeq sequencer at Swift Biosciences. Additional libraries for 12 samples of North Atlantic harbor porpoises were prepared at BGI Inc. using their proprietary protocol, indexed and pooled for paired-end 150 bps sequencing on one lane of HiSeq-4000 at BGI Inc. The total sequencing effort produced reads for 56 individuals (Table S2). Previously published reads from one additional finless porpoise sequenced with a Hiseq-2000 (Yim et al., 2014) were retrieved from NCBI (Table S2). For this individual, we down-sampled the raw FASTQ files to extract 0.5% of the total reads and used the 5,214,672 first reads to assemble the mitogenomes.

### 2.2. Data cleaning

The quality of the reads was first evaluated for each sample using *FastQC* v.0.11.5 (Andrews, 2010). *Trimmomatic* v.0.36 (Bolger et al., 2014) was used to remove low quality regions, overrepresented sequences and Illumina adapters. Different filters were applied according to the type of Illumina platform used (see Text S1 for details). Only mated reads that passed *Trimmomatic* quality filters were used for the subsequent analyses.

### 2.3. Mitogenome assembly

Porpoises mitogenomes assemblies were reconstructed using two different approaches. First, we used *Geneious* v.8.1.8 (Kearse et al., 2012) to perform a direct read mapping to the reference mitogenome of the harbor porpoise (accession number AJ554063; Arnason et al., 2004), with a minimum mapping quality of 20 (see Fig. S1 for details). This step was followed by a reconstruction of the consensus sequences (Table S2). The second approach implemented in *MITOBIM* (Hahn et al., 2013) is an hybrid approach combining a baiting and iterative elongation procedure to perform a *de-novo* assembly of the mitogenome (see details in Text S2). We visually compared the assemblies provided by the two methods in *Geneious* to assess and resolve inconsistencies (Text S2 and Table S2).

In addition to the 57 assembled individuals, we retrieved six porpoise mitogenomes from Genbank (Table S2). We also added eight complete mitogenomes from four outgroup species: one narwhal (*Monodon monoceros*) (Arnason et al., 2004), three bottlenose dolphins *(Tursiops truncatus*) (Moura et al., 2013), one Burrunan dolphin (*Tursiops australis*) (Moura et al., 2013) and three orcas (*Orcinus orca*) (Morin et al., 2010).

### 2.4. Sequences alignments

We performed the alignment of the 71 mitogenomes with *MUSCLE* (Edgar, 2004) using default settings in *Geneious*. A highly repetitive region of 226 bps in the D-loop was excluded from the final alignment (from position 15,508 to 15,734) because it was poorly assembled, and included many gaps and ambiguities. We manually annotated the protein-coding genes (CDS), tRNA and rRNA of the final alignment based on a published harbor porpoise mitogenome (Arnason et al., 2004). Contrary to the remaining CDSs, ND6 is transcribed from the opposite strand (Clayton, 2000). Therefore, to assign the codon positions in this gene, we wrote a custom script to reverse complement ND6 in the inputs of all the analyses that separates coding and non-coding regions of the mitogenomes. This led to a 17 bps longer alignment due to the overlapping position of ND5 and ND6.

### 2.5. Phylogenetic relationships

We estimated the phylogenetic relationships among the assembled mtDNA sequences using 3 approaches: a maximum-likelihood method in *PHYML* v3.0 (Guindon et al., 2010) (ML); a distance based method using the Neighbour-Joining algorithm (NJ) in *Geneious*; and an unconstrained branch length Bayesian phylogenetic tree (BI) in *MrBayes* v3.2.6 (Ronquist et al., 2012). We used the Bayesian information criterion (BIC) to select the substitution model that best fits the data in *jModelTest2* v2.1.10 (Darriba et al., 2012). The best substitution model and parameters were used in the ML, NJ and BI approaches. For the ML approach, we fixed the proportion of invariable sites and the gamma distribution parameters to the values estimated by *jModelTest2.* The robustness of the ML and NJ tree at each node was assessed using 10,000 bootstrap replicates. For the Bayesian inference, a total of 1×10^6^ MCMC iterations was run after a burn-in of 1×10^5^ steps, recording values with a thinning of 2,000. We performed 10 independent replicates and checked the consistency of the results. Runs were considered to have properly explored the parameter space if the effective sample sizes (ESS) for each parameters was greater than 200 and by visually inspecting the trace plot for each parameter using *Tracer* v1.6 (Rambaut et al., 2014). We assessed the statistical robustness and the reliability of the Bayesian tree topology using the posterior probability at each node.

Phylogenetic inference can be influenced by the way the data is partitioned and by the substitution model used (Kainer and Lanfear, 2015). Therefore, in the ML analysis, besides trying to find the best substitution model for the complete mitogenome using *jModelTest2,* we also tested a partitioned scheme of the mitogenomes into 42 subsets using *PARTITIONFINDER* v1.1.0 (Lanfear et al., 2012) and search for the optimal substitution model for each partition defined. The partition scheme included (1) one partition per codon position for each CDS (3×13=39 partitions in total); (2) one partition for the non-coding regions; (3) one partition for all rRNAs genes; and (4) one partition for all tRNAs genes. We used the greedy algorithm of *PARTITIONFINDER*, the *RAxML* set of substitution models and the BIC to compare the fit of the different models. ML tree was reconstructed using the results of *PARTITIONFINDER* in *RAxML* v8 (Stamatakis, 2014). The robustness of partitioned ML tree was assessed using 10,000 rapid bootstraps (Stamatakis et al., 2008).

Finally, the four phylogenetic trees were rooted with the eight outgroup sequences and plotted using the R package *ggtree* v1.4 (Yu et al., 2018).

### 2.6. Divergence time estimate

We estimated the divergence time of the different lineages using a time-calibrated Bayesian phylogenetic analysis implemented in *BEAST* v2.4.3 (Bouckaert et al., 2014). We assumed two calibration points in the tree: (1) the split between the Monodontidae and Phocoenidae, node calibrated with a lognormal mean age at 2.74 Myr (McGowen et al., 2009) (sd=0.15) as a prior and (2) the split between the Pacific and Atlantic harbor porpoise lineages, node calibrated with a Uniform distribution between 0.7 and 0.9 Myr as a prior (Tolley and Rosel, 2006).

Divergence times were estimated using a relaxed log-normal molecular clock model for which we set the parameters *ucldMean* and *ucldStdev* to exponential distributions with a mean of 1 and 0.3337, respectively. We used a Yule speciation tree model and fixed the mean of the speciation rate parameter to 1. The *BIC* was used in *jModelTest2* to identify the substitution model best fitting to the data, using the empirical base frequencies. We assumed a substitution rate of 5×10^-8^ substitutions per-site and per-year (Nabholz et al., 2007). A total of 1.2×10^9^ MCMC iterations were run after a burn-in length of 1.2×10^8^ iterations, with a thinning of 5,000 iterations. We performed eight independent replicates and checked for the consistency of the results among replicates. A run was considered as having converged if the ESS values were greater than 200, and if they produced consistent trace plots using *Tracer* v1.6. Subsequently, we combined all runs together after a burn-in of 98% using *LogCombiner* (Bouckaert et al., 2014). The best supported tree topology was the one with the highest product of clade posterior probabilities across all nodes (maximum clade credibility tree), estimated using *TreeAnnotator* (Bouckaert et al., 2014). We also calculated the 95% highest posterior density for the node ages using *TreeAnnotator*. The final chronogram was rooted with the eight outgroups sequences and plotted using *FigTree* v.1.4.3 (Rambaut and Drummond, 2012).

### 2.7. Genetic diversity within species and subspecies

We subdivided each species into their distinct lineages in order to compare genetic diversity at the different taxonomic level. Specifically, we divided the harbor porpoise into five subgroups, North Pacific (*P. p. vomerina*), Black Sea (*P. p. relicta*), Mauritanian, Iberian (*P.p. meridionalis*) and North Atlantic (*P. p. phocoena*) in accordance with the subdivisions proposed for this species in the literature (Fontaine, 2016). Finless porpoise was split into Indo-Pacific finless (*N. phocaenoides;* IPF), East Asian finless (*N. a. sunameri*; EAF) and Yangtze finless porpoises (*N. a. asiaeorientalis*; YFP). Additionally, we subdivided the other groups into lineages that were as divergent or more divergent than the sub-species that were described in the literature. This includes splitting the North Pacific harbor porpoises into NP1 and NP2, Dall’s porpoises into DP1 and DP2 and spectacled porpoises into SP1 and SP2 to reflect the deep divergence observed in the phylogenetic tree within these three lineages (Fig. 1b and S2).

For each species and subgroup, several summary statistics capturing different aspects of the genetic diversity were calculated for different partition of the mitogenome, including the whole mitogenomes, the non-coding regions (i.e. inter-gene regions and D-loop) and the CDSs (i.e. 13 protein coding genes excluding tRNAs and rRNA). The number of polymorphic sites, nucleotide diversity (*π*), number of singletons, number of shared polymorphisms, number of haplotypes, haplotype diversity and Watterson estimator of *θ* were calculated. For CDSs, we also estimated the number of synonymous (*#Syn*) and non-synonymous mutations (*#NSyn*), *π* based on synonymous (*π_S_*) and non-synonymous mutations (*π_N_*), and the ratio *π_N_/π_S_*. All these statistics were computed in *DnaSP* v.5.10.01 (Librado and Rozas, 2009). Since we only have a unique sample for IPF, DP1 and SP1 we did not estimate these statistics for these lineages.

Differences in sample sizes can potentially influence some of these statistics. As our sample size ranged from three to 26 individuals per group, we used a rarefaction technique (Sanders, 1968) to account for the differences in sample size. We assumed a sample size of three individuals in order to compare the genetic diversity among lineages that have different sample sizes. For each lineage, we randomly subsampled 2,500 times a pool of three sequences and estimated the median, mean and 95% confidence interval for *π*.

### 2.8. Test for selective neutrality

We tested for evidence of natural selection acting on the mitogenomes using a McDonald-Kreitman test (MK-tests) (McDonald and Kreitman, 1991). This test infers deviation from the neutral theory by comparing the ratio of divergence between species (*d_N_/d_S_*) *versus* polymorphism within species (*π_N_/π_S_*) at synonymous (silent) and non-synonymous (amino acid-changing) sites in protein coding regions using the so-called neutrality index (*NI*). *NI* is 1 when evolution is neutral, greater than 1 under purifying selection, and less than 1 in the case of adaptation. MK-tests were conducted on the 13 CDSs regions of the mitogenome using *DnaSP*. We conducted this test in two different ways: first comparing all the interspecific lineages to a same outgroup, the killer whale, and second comparing all interspecific lineages to each other in order to assess how the MK-tests were affected by the outgroup choice. The significance of the *NI* values was evaluated using a G-tests in *DnaSP*. Furthermore, the distribution of *NI* values for each lineage were compared among each other using a PERMANOVA with the R package *RVAideMemoire* (Hervé, 2019). Pairwise comparisons were assessed with a permutation tests and were adjusted for multiple comparisons using the false rate discovery method (Benjamini and Hochberg, 1995). The PERMANOVA and pairwise comparisons were conducted using 9999 permutations. The neutral theory predicts that the efficacy of purifying selection increases with *Ne* (Kimura, 1983). Under these assumptions, *Ne* is expected to be proportional to *NI* (Hughes, 2008; Phifer-Rixey et al., 2012). To test this hypothesis, we assessed the correlation between *NI*s and *π* derived by rarefaction as a proxy of *Ne*.

In addition to the MK-tests, we quantified the branch-specific non-synonymous to synonymous substitution ratios (*d_N_/d_S_*) to infer direction and magnitude of natural selection along porpoise phylogenetic tree. To estimate the branch-specific ratio we first counted the number of synonymous (*#S*) and non-synonymous *(#NS*) substitutions for the 13 CDSs. Then *#S* and *#NS* were mapped onto a tree using the mapping procedure of Romiguier et al. (2012). Next, we divided *#S* and *#N* by the number of synonymous and nonsynonymous sites to obtain an approximation of *d_S_*and *d_N_*, respectively. More specifically, we first fitted the YN98 codon model using the *BPPML* program (Dutheil and Boussau, 2008), then we mapped the estimated *d_N_*/*d_S_* values onto the branches of the ML tree using the program *MAPNH* of the *TESTNH* package v1.3.0 (Dutheil et al., 2012). Extreme *d_N_*/*d_S_* ratio (> 3) are often due to branches with very few substitution (*d_N_* or *d_S_*) (Romiguier et al., 2012) and were discarded. We compared the distribution of *d_N_*/*d_S_* among species (*i.e.*, across all the branches) using a PERMANOVA. Finally, the estimated ratios were correlated with *π* obtained by rarefaction using a Pearson’s correlation tests in R (R Core Team, 2018). To do so, we pooled the signal from each lineage as a single data point as suggest by Figuet et al. (2014). We considered the intraspecific and interspecific lineages, except for those where no non-synonymous substitutions were observed (ex. NP2). Within a lineage, *π* was summarized as the mean of the log_10_-transformed value of its representatives and the *d_N_*/*d_S_* was obtained by summing the non-synonymous and synonymous substitution counts across its branches and calculating the ratio (Figuet et al., 2014).

### 2.9. Inference of demographic changes

To conduct reliable demographic inferences, we investigated changes in *Ne* through time for the lineages that included a sample size ≥ 10 to conduct reliable demographic inferences. This includes the vaquitas and North Pacific harbor porpoise lineage NP1. We first tested for deviations from neutral model expectations using three statistics indicative of historical population size changes: Tajima’s *D* (Tajima, 1989), Fu and Li’s *D\** and *F\** (Fu and Li, 1993) in *DnaSP*. The *p*-values were assessed using coalescent simulations in *DnaSP* using a two tailed test as described in Tajima (1989). We then reconstructed the mismatch distributions indicative of population size changes using *Arlequin* v.3.5.2.2 (Excoffier and Lischer, 2010). Mismatch distribution were generated under a constant population size model and a sudden growth/decline population model (Schneider and Excoffier, 1999). This later model is based on three parameters: the population genetic diversity before the population change (*θ_i_*); the population genetic diversity after the population change (*θ_f_*), and the date of the change of population size in mutational units (*τ* = *2.µ.t*, where *µ* is the mutation rate per sequence and generation and *t* is the time in generations). These parameters were estimated in *Arlequin* using a generalized non-linear least-square approach. The 95% confidence interval was obtained using 10,000 parametric bootstraps (Schneider and Excoffier, 1999). Finally, we used the coalescence based Bayesian Skyline Plot (BSP) (Drummond et al., 2005) to estimates demographic changes in *Ne* back to the *T_MRCA_*. BSP analysis was performed in *BEAST* v2.4.3 using the empirical bases frequencies and a strict molecular clock. We applied *jModelTest2* separately to both lineages to evaluate the best substitution models. We assumed a substitution rate of 5×10^-8^ substitutions per site and per year (Nabholz et al., 2007) in order to obtain the time estimates in years. We conducted a total of 10 independent runs of 10^8^ MCMC iterations following a burn-in of 1×10^7^ iterations, logging every 3000 iterations. We constrained *Ne* between 0 and 150,000 individuals and between 0 and 380,000 individuals for the vaquita and the NP1 harbor porpoise lineage, respectively. This upper boundary on *Ne* was empirically set to encompass the entire marginal posterior distribution. All other parameters were kept at default values. The convergence of the analysis was assessed by checking the consistency of the results over 10 independent runs. For each run, we also used *Tracer* to inspect the trace plots for each parameter to assess the mixing properties of the MCMCs and estimate the ESS value for each parameter. Run were considered as having converged if they displayed good mixing properties and if the ESS values for all parameters was greater than 200. We discarded the first 10% of the MCMC steps as a burn-in and obtained the marginal posterior parameter distributions from the remaining steps using *Tracer*. *Ne* values were obtained by assuming a generation time of 10 years (Birkun and Frantzis, 2008).

## 3. Results

### 3.1. Porpoise mitogenomes assemblies

A total of 57 mitogenomes of the seven species of porpoise (Table S1) were newly sequenced and assembled using Illumina sequencing. After read quality check, trimming, and filtering (Table 1 and S2), between 1,726,422 and 67,442,586 cleaned reads per sample were used to assemble the mitogenomes. The two methods used to assemble the mitogenomes delivered consistent assemblies with an average depth coverage ranging from 15 to 4,371X for *Geneious* and 17 to 4,360X for *MITOBIM* (Table 1 and S2). In total, 35 of the 57 mitogenome assemblies were strictly identical with both methods. The 22 remaining assemblies included between 1 and 4 inconsistencies which were visually inspected and resolved (Text S2 and Table S2). Augmented with the 14 previously published mitogenome sequences, the final alignment counted 71 mitogenome sequences of 16,302 bps and included less than 0.2% of missing data. The alignment was composed of 627 singletons and 3,937 parsimony informative sites defining 68 haplotypes (including the outgroup sequences). Within the 63 ingroup porpoise sequences, we observed 2,947 segregating sites, including 242 singletons and 2,705 shared polymorphisms defining 58 distinct haplotypes with a haplotype diversity of 99.6%.

### 3.2. Phylogenetic history of the porpoises

A Hasegawa-Kishino-Yano (HKY+G, Gamma=0.19) model was selected as the best-fitting nucleotide substitution model. Phylogenetic inferences using a maximum-likelihood (ML) approach based on the whole mitogenome (Fig. 1b and Fig. S2a) or on a partitioned data set (Fig. S2b), a distance-based neighbor-joining method (Fig. S2c), and a Bayesian approach (Fig. S2d) all produced concordant phylogenies (i.e., similar topologies and statistical supports). All phylogenies were fully resolved with high statistical support at the inter- and intra-specific levels (bootstrap supports ≥93% or node posterior probability of one). One exception was the node 5 grouping the Burmeister’s and spectacled porpoises in the neighbor-joining tree (Fig. S2b) as it displayed a slightly lower bootstrap support of 61% (red star in Fig. 1b and S2b), but it was fully supported by the other approaches (Fig. 1b and S2).

The resulting phylogenetic reconstruction (Fig. 1b) showed that all porpoises formed a monophyletic group (node 1). The most basal divergence in the tree split the more tropical finless porpoises from the other temperate to subpolar porpoises. Then, the temperate species split into two clades (node 2) composed of two reciprocally monophyletic groups. The first is composed of the southern hemisphere species (spectacled and Burmeister’s porpoises) and vaquitas (node 3). The second is composed of the porpoises from the northern hemisphere (harbor and Dall’s porpoises, node 4). In contrast with a previous phylogenetic study based on control region sequences (Rosel et al., 1995b), the phylogenetic tree based on the whole mitogenome suggested that vaquitas split from a common ancestor to the spectacled and Burmeister’s porpoises (node 3), and thus that the two species from the southern hemisphere are more closely related to each other (node 5) than they are to vaquitas. Finally, the mitogenome tree supported the monophyly of each recognized species (nodes 6-11).

Intraspecific subdivisions were also evident from the mitogenome phylogeny in some species, such as in the harbor and finless porpoises (Fig 1b). In the harbor porpoises, the split between the North Atlantic and North Pacific subspecies constituted the deepest divergence of all intraspecific subdivisions across all species. Within the North Atlantic harbor porpoises, further subdivisions were also observed and corresponded to the three previously described ecotypes or sub-species (Fontaine, 2016). These included the relict population in the Black Sea, the harbor porpoises from the upwelling waters with two closely related but distinct lineages in Iberia and Mauritania, and the continental shelf porpoises further north in the North Atlantic. Finally, within in the North Pacific, two cryptic subgroups were also observed (NP1 and NP2; Fig. 1b). Among the finless porpoises, the three taxonomic groups currently recognized (Zhou et al., 2018), including IPF and the two closely related species of narrow-ridged finless porpoises, were clearly distinct from each other on the mitogenome phylogenetic tree (node 11). Finally, despite a limited sampling, Dall’s (node 7) and spectacled porpoises (node 10) each also displayed distinct lineages (DP1/DP2 and SP1/SP2, respectively; Fig. 1b and S2) as divergent as those observed in the harbor and finless porpoises.

The time-calibrated Bayesian mitochondrial phylogeny (Fig. 2 and Table S5) suggested that all extant porpoises find their common ancestor ∼5.42M years ago (95% Highest Posterior Density, HPD, 4.24-6.89; node 1). This time corresponds to the split between the finless and the other porpoise species. Spectacled, vaquita and Burmeister’s porpoises diverged from harbor and Dall’s porpoise ∼4.06 Myr ago (95% HPD, 3.15-5.12; node 2). The split between vaquitas, spectacled and Burmeister’s porpoises was estimated at ∼2.39 Myr (95% HPD, 1.74-3.19; node 3) and between spectacled and Burmeister’s at ∼2.14 Myr (95% HPD, 1.51-2.86; node 4). The Dall’s and harbor porpoises split from each other ∼3.12 Myr ago (95% HPD, 2.31-3.98; node 5). Finally, the common ancestor of the subdivisions observed within each species was dated within the last million years (nodes 6-11).

**Fig. 2.**
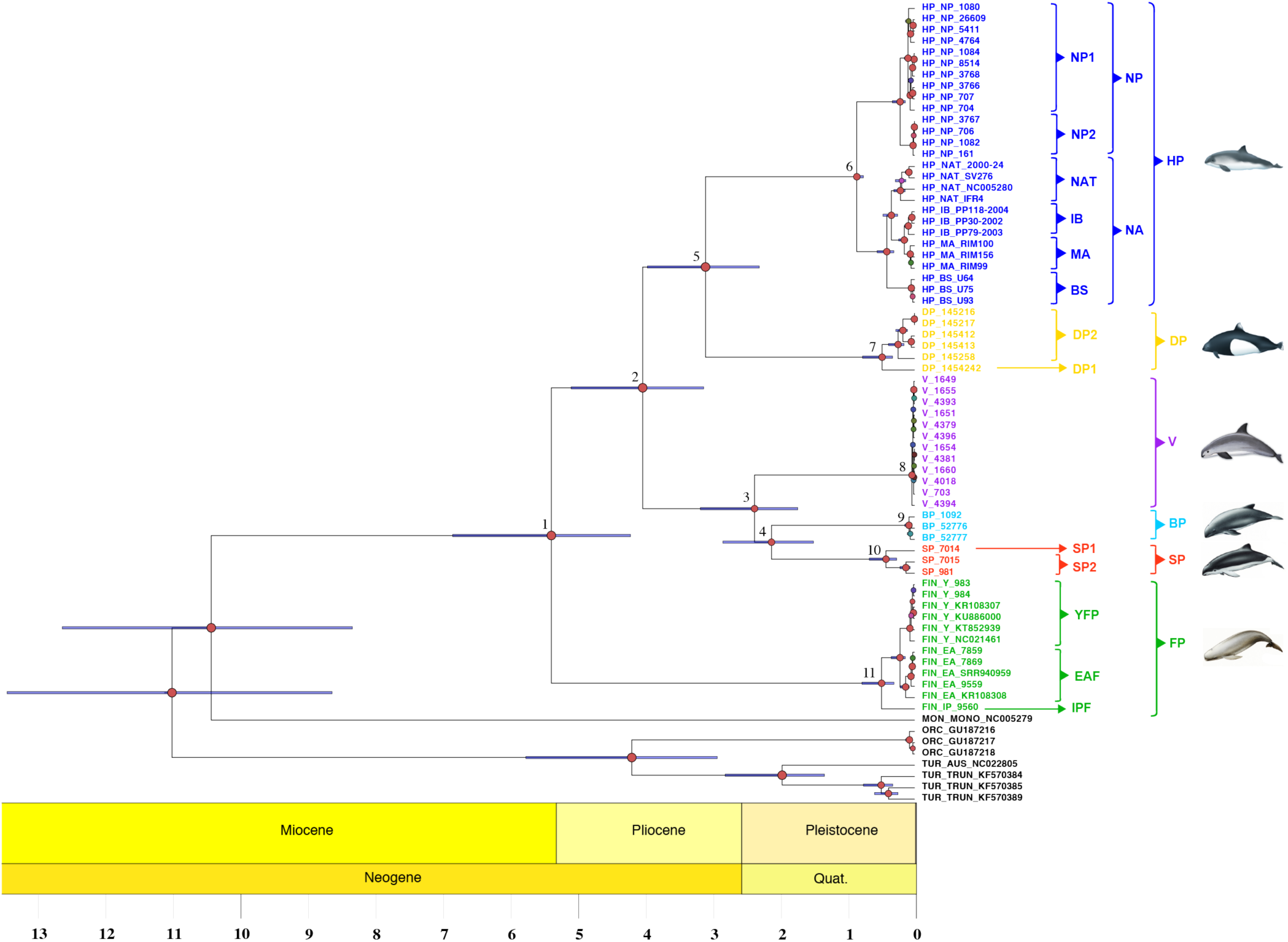
Bayesian chronogram of the porpoise family. The tree represents the maximum clade credibility tree. Red node labels indicate posterior probabilities of 1; node position indicates median node age estimates and the error bars around the node indicate the 95% highest posterior density of the estimated node age. Time is in millions of years. Numbers at the nodes are discussed in the text. The acronyms are provided in Table 1.

#### Genetic diversity of the mitogenome

Mitochondrial genetic diversity varied greatly among species (Table 2 and Fig. 3). The highest values of *π* were observed in the harbor porpoises (*π*=1.15%), followed by the spectacled (*π*=0.60%), Dall’s (*π*=0.50%), finless (*π*=0.35%), and Burmeister’s porpoises (*π*=0.13%), while vaquitas displayed the lowest values (*π*=0.02%). The variation among species was strongly related to the occurrence of distinct mitochondrial lineages within species that corresponds to ecologically and genetically distinct sub-species or ecotypes (Table 2 and Fig. 3). Once the lineages that included more than three sequences were compared to each other while accounting for the difference in sample size using a rarefaction procedure (Sanders, 1968) (Fig. 3), π values were more homogeneous among lineages and species, with however some variation. The most diversified lineages included DP2 in Dall’s porpoise (*π*=0.32%) and the North Atlantic (NAT) lineage in harbor porpoise (*π*=0.33%). In contrast, the vaquita lineage (*π*=0.02%), harbor porpoises North Pacific lineages NP2 (*π*=0.02%) and Black Sea (BS) (*π*=0.07%), and the Yangtze finless porpoise YFP lineage (*π*=0.06%) displayed the lowest nucleotide diversity. The other lineages displayed intermediate *π* values.

**Fig. 3.**
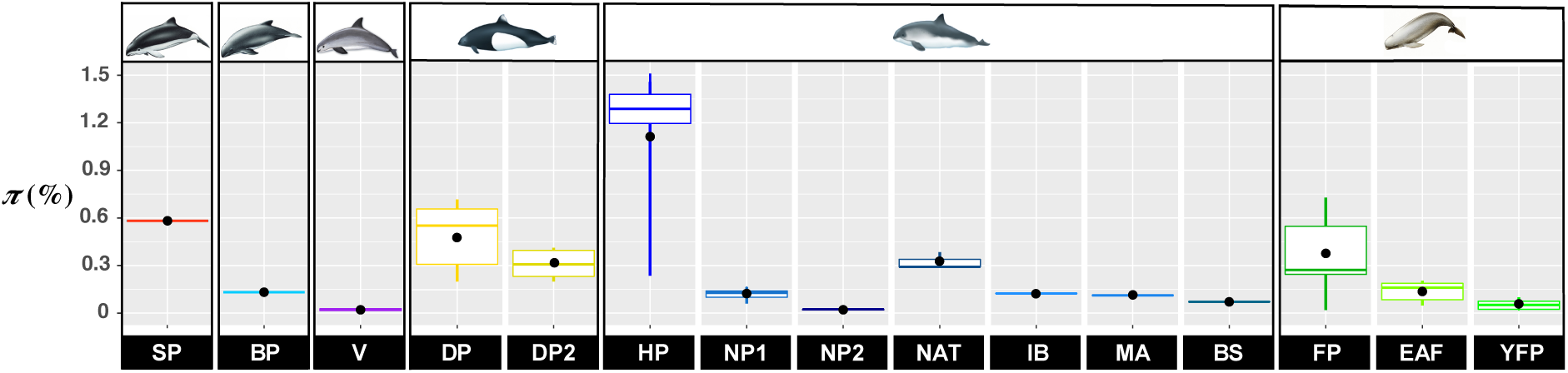
Nucleotide diversity (*π*) among groups and species of porpoises. The median and mean *π* values are represented respectively by the colored line in the boxplot and the black dot. The whiskers represent the 95% confidence interval. The boxes represent the upper and lower quartile. The acronyms are provided in Table 1.

**Table 2.**
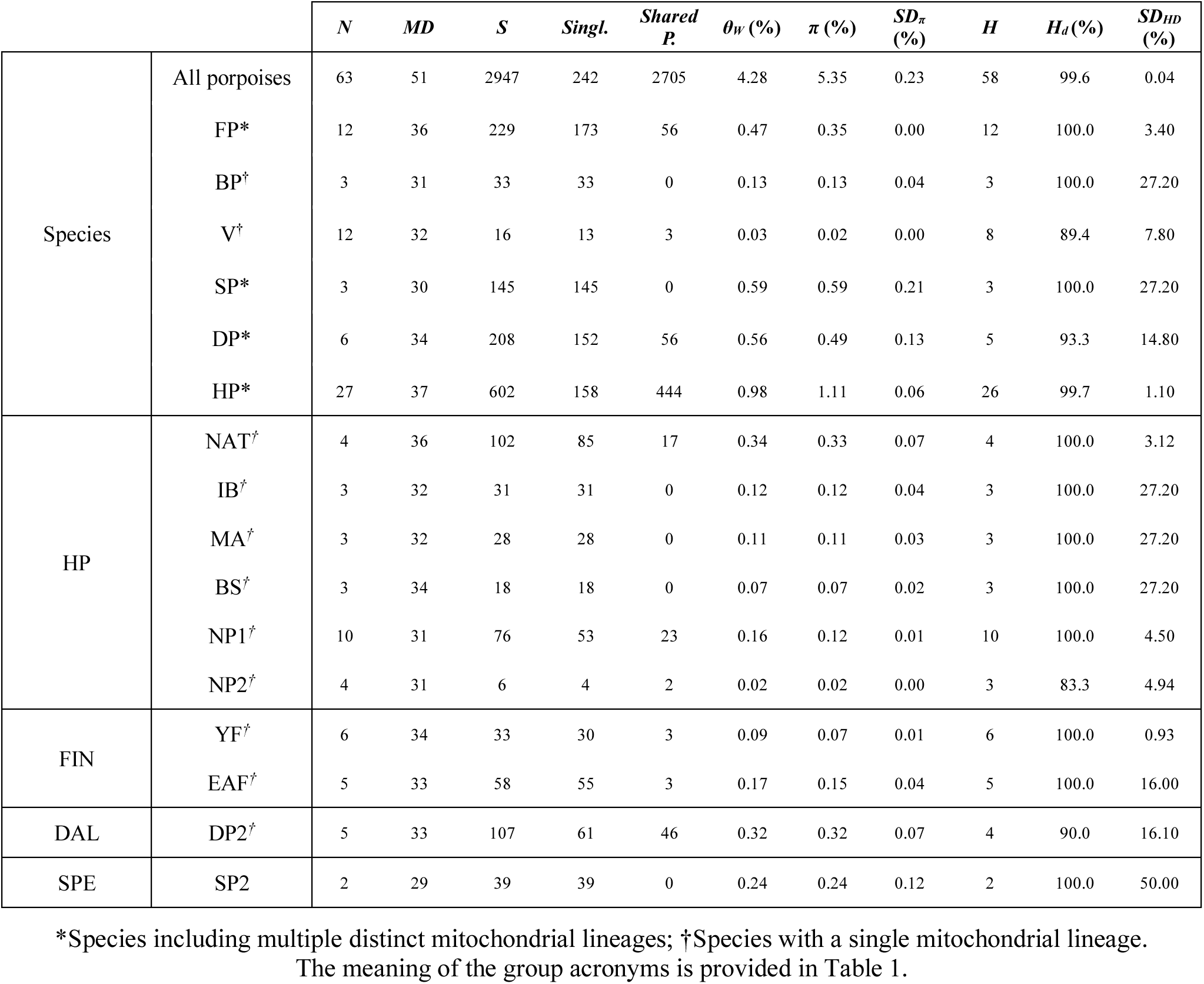
Summary statistics describing the genetic diversity of the mitochondrial genomes among porpoise species and their distinct lineages. The statistics includes the sample size (*N*), number of sites with missing data (*MD*), segregating sites (*S*), singletons (*Singl*.), shared polymorphism (*Shared P.),* theta from *S (θ_W_*), average nucleotide diversity per site (*π*) and its standard deviation (*SD_π_*), number of haplotypes (*H*), haplotypic diversity (*H_d_*) and its standard deviation (*SD_HD_*).

### 3.3. Molecular evolution of the mitogenome

The nucleotide diversity also varied greatly along the mitogenome. It was lowest in the origin of replication, tRNA and rRNAs, intermediate in the coding regions and highest in non-coding regions (Fig. S3). This result indicates different levels of molecular constraints along the mitogenome.

The *π_N_/π_S_* ratio in the 13 CDSs, displaying the relative proportion of non-synonymous *versus* synonymous nucleotide diversity, was lower than one in all the lineages. This is consistent with purifying selection acting on the coding regions. At the species level, the ratio *π_N_/π_S_* ranged from 0.04 in Dall’s and spectacled to 0.10 in finless porpoises (Table S3). The vaquita displayed an intermediary value of 0.06. Within each species, *π_N_/π_S_* also varied markedly across lineages: in the harbor porpoise, *π_N_/π_S_* ratios ranged from 0 in the North Pacific NP2 lineage to 0.21 in the Black Sea BS lineage; in the finless porpoises from 0.14 in EAF to 0.17 in YFP; 0.05 in the DP2 Dall’s porpoise lineage; and 0.06 in SP2 spectacled porpoise lineage (Table S3).

The branch-specific non-synonymous to synonymous substitution rate (*d_N_/d_S_*, Fig. S4a) were fairly conserved across the phylogenetic tree and ranged from 0 in the finless porpoise to 1.08 in the harbor porpoise, with a median value at 0.12. A *d_N_/d_S_* lower than one implies purifying selection. Thus, similar to *π_N_/π_S_*, the branch-specific *d_N_/d_S_* suggested that the porpoise mitogenomes were mostly influenced by purifying selection. Furthermore, the *d_N_/d_S_* ratios did not differ significantly among species (PERMANOVA, *p*-value=0.49). Interestingly, the *d_N_/d_S_* ratio was negatively correlated with the nucleotide diversity (Fig. S4b; Pearson’s *r* = −0.64, *p*-value = 0.01) suggesting that purifying selection removes deleterious mutations more effectively in genetically more diversified lineages.

The Mcdonald-Kreitman (MK) tests using first the orca as an outgroup showed that all the lineages for each species had neutrality index (*NI)* values greater or equal to one (Fig. 4a). In particular, some lineages displayed *NI* values significantly higher than one (G-tests, *p*-value<0.05), consistent with a signal of purifying selection. These included the EAF and YFP lineages in the finless porpoises and the MA, IB and BS in the harbor porpoises (Fig. 4a). Vaquitas and NP2 harbor porpoise lineages also displayed marginally significant *NI* values (*NI* > 2, *p*-value≤0.10; Fig. 4a). The remaining lineages showed *NI*s close to one suggestive of selective neutrality. The MK tests applied to all pairs of interspecific lineages showed *NI* values often higher than one (Fig. 4b and Fig. S5a). The values were especially high (Fig. S5a) and significant (Fig. S5b) when comparing the harbor porpoise lineages (MA, IB, and BS) with the finless porpoise linages (YFP and EAF). The variation in the distribution of *NI* among interspecific lineages (Fig. 4b) showed that these same lineages displayed significantly larger *NI* values compared to spectacled SP2 and Dall’s DP2 porpoise lineages (PERMANOVA, *p*-value<0.001 and all pairwise comparisons have a *p*-value<0.04 after False Rate discovery adjustment). We observed a significant negative correlation between *π* and *NI* (Pearson’s *r* = −0.28, *p*-value=0.003), suggesting that purifying selection is stronger in lineages with small *Ne* or that demographic events impacted the polymorphism of these lineages.

**Fig. 4.**
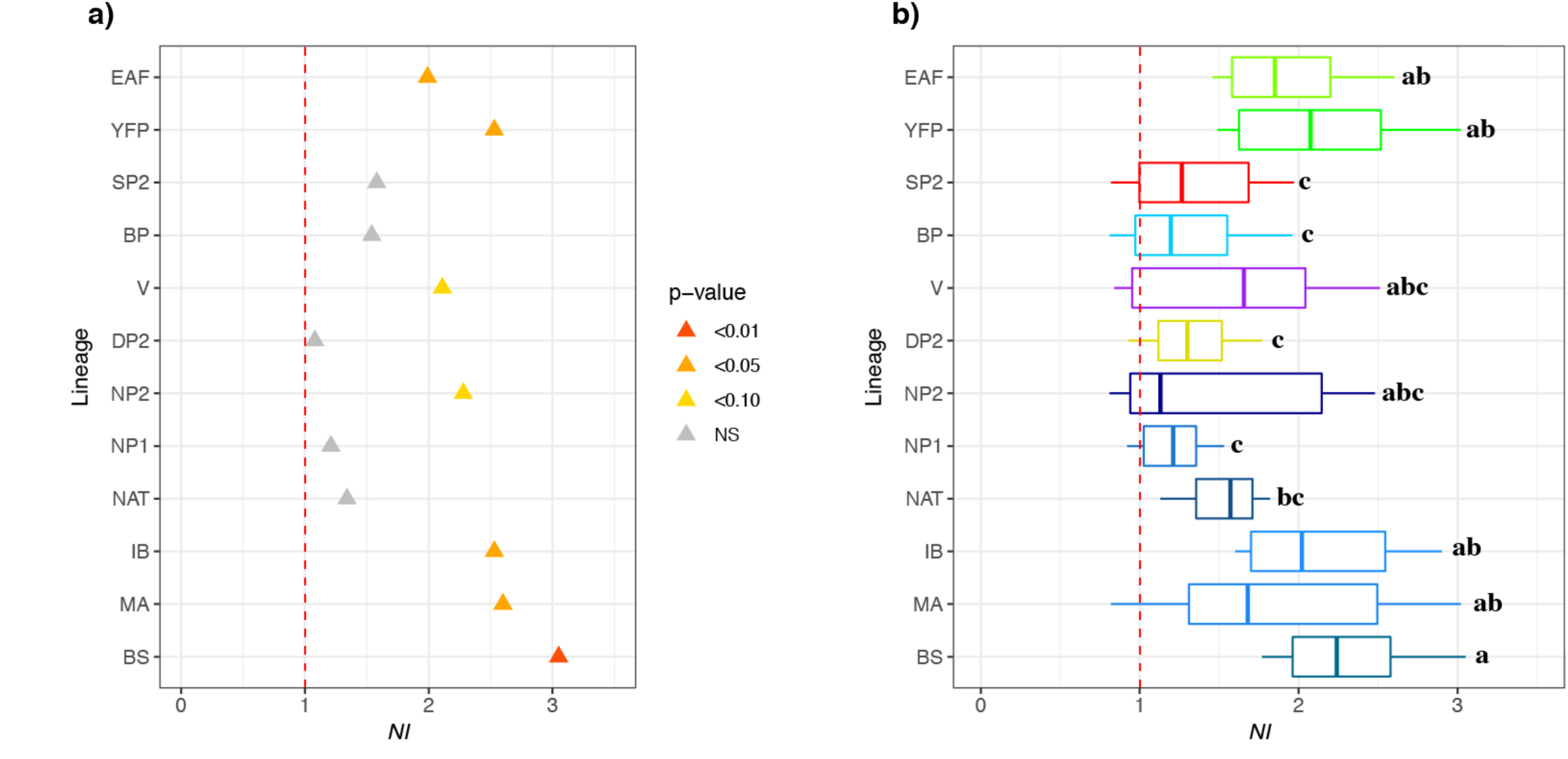
Mcdonald-Kreitman tests on the 13 protein coding regions of the mitogenome among porpoise subgroups. (a) Neutrality index (*NI*) estimated using the orca as outgroup. (b) NI distributions per mitochondrial lineages, calculated using various outgroups, including orca and all possible interspecific comparisons. The letters on the right of the boxplots indicate significant differences in the mean *NI* between the different lineages (i.e. boxplot with different letters are statistically different from one another). The red dashed lines represent the limit at which *NI* reflects positive (*NI*<1), neutral (*NI* =1), or purifying (*NI*>1) selection. NS: Not Significant. The acronyms are provided in Table 1.

### 3.4. Demographic history

The vaquita displayed significant departure from neutral constant population size expectations with significant negative values for Fu and Li’s *D\** and *F\**, but not Tajima’s *D* (Table 3). This result indicates a significant excess of singleton mutations compare to a neutral expectation, consistent with a bottleneck or a selective sweep. In contrast, the harbor porpoise NP1 lineage did not show any such deviation, even though all the statistics displayed negative values.

**Table 3.** Neutrality tests based on the site frequency spectrum. The neutrality tests were only applied to lineages where at the sample size (*n)* was at least 10. The statistics includes the Tajima’s *D*, Fu and Li’s *D\**, and Fu and Li’s *F*.

| Lineage | $D$ | $D^*$ | $F$ |
| --- | --- | --- | --- |
| V ($n=12$) | -1.44<sup>NS</sup> | -1.89* | -1.84* |
| NP1 ($n=10$) | -1.18<sup>NS</sup> | -1.27<sup>NS</sup> | -1.28<sup>NS</sup> |

NS, Not significant; \*, p-value < 0.05. The acronyms are provided in Table 1.

The mismatch distribution and the coalescent-based BSP also captured this contrast (Fig. 5). For the North Pacific harbor porpoise NP1 lineage, the mismatch distribution was consistent with an ancient population expansion (Fig. 5a) with a modal value close to 20 differences on average between pairs of sequences. Despite the ragged distribution and large 95% CI, the best fitted model suggested an ancient increase in genetic diversity (*θ=2.Ne.µ*) by a factor of 40 after a period (*τ=2.t.µ)* of 18 units of mutational time. This old expansion was also detected by the BSP analysis (Fig. 5c). Indeed, NP1 displayed an old steady increase in *Ne* with time since the most recent common ancestor (*T_MRCA_*) 16,166 years before present (yrs. BP) (95%CI: 12,080–20,743), with a median *Ne* increasing from 1,660 to 6,436 (Fig. 5c). For the vaquita lineage, the mismatch distribution and the BSP analyses both supported a much more recent expansion than in NP1. The mismatch distribution (Fig. 5b) showed an increase of *θ* by a factor of 1,000 after a *τ* of four units of mutational time. The mode of the bell shape distribution for the best fitted model was around three differences among pairs of sequences, which is consistent with a recent population expansion. The BSP analysis (Fig. 5d) detected this expansion and dated it back to 3,079 yrs. BP (95%CI: 1,409–5,225), with median *Ne* increasing from 613 (95%CI: 45-5,319) to 1,665 (95%CI: 276-9,611). Thus, the estimated current *Ne* was three to six times lower than in NP1 (Fig. 5d).

**Fig. 5.**
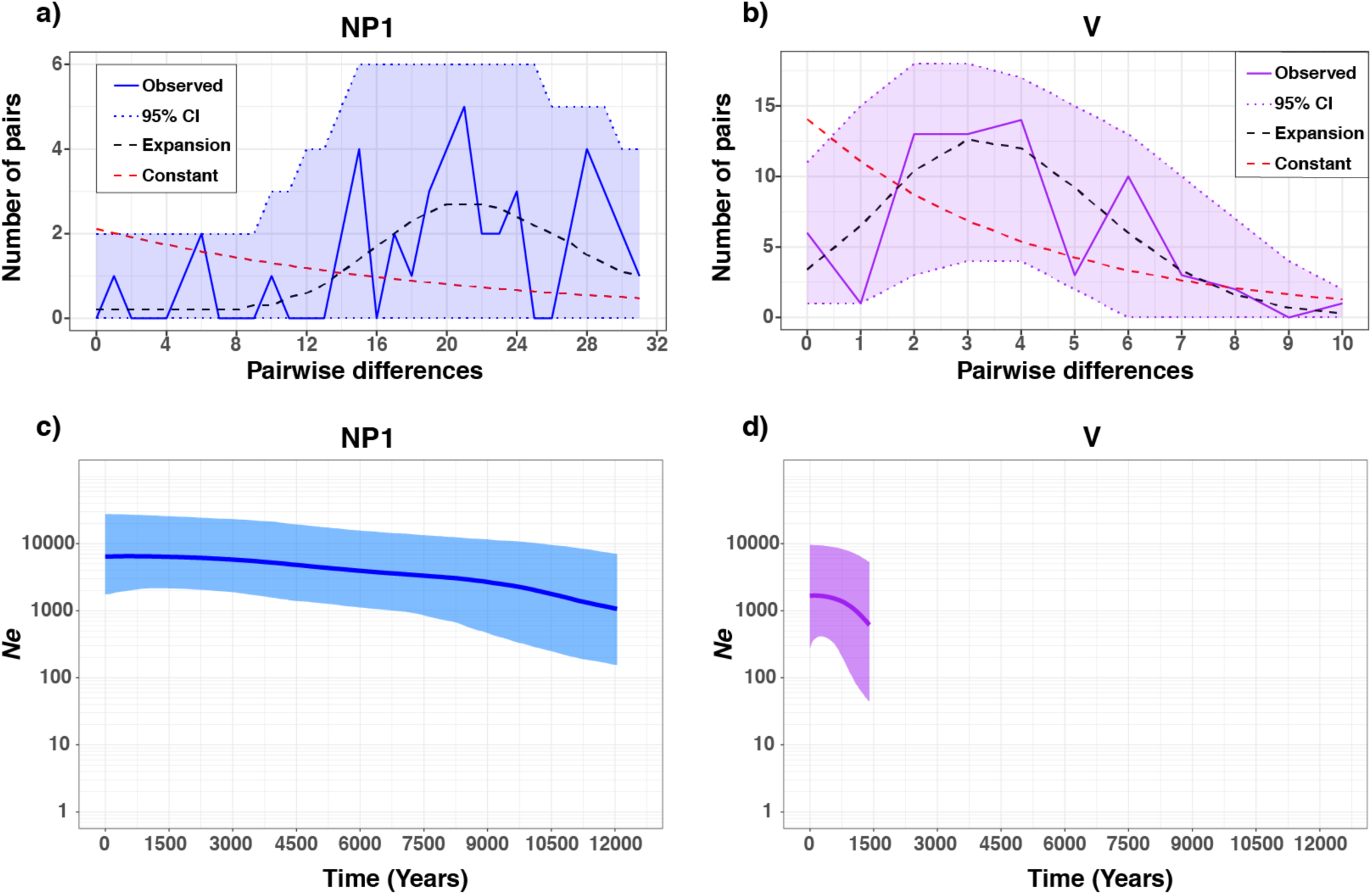
Mismatch distributions (a and b) and Bayesian Skyline Plots (c and d) for the North Pacific harbor porpoise NP1 lineage and for the vaquita. The Skyline Plots (c and d) show the temporal changes in mitochondrial diversity. The y-axis of the Bayesian Skyline Plot shows the genetic diversity expressed as the effective female population size (*Ne*). The bold line inside each Bayesian Skyline Plot represents the median value and thin lines the 95% HPD intervals. The acronyms are provided in Table 1.

## 4. Discussion

The phylogeny of the Phocoenidae has been debated for decades, in part due to the lack of polymorphism and statistical power that came from the analyses of short fragments of the mitochondrial genome (McGowen et al., 2009; Rosel et al., 1995b). Using massive parallel sequences technologies, the analyses of newly sequenced and assembled whole mitogenomes from all the species and subspecies of porpoises provide the first robust comprehensive picture of the evolutionary history of the porpoises. The phylogenetic relationships estimated here delivered a fully resolved evolutionary tree (Fig 1b and S2). While most of the phylogenetic relationships were suggested previously (McGowen et al., 2009; Rosel et al., 1995b), the resolution and statistical support recovered here was maximal. Our results support the monophyly and branching of each species and subspecies. Moreover, the comparative view of the mitochondrial polymorphism within and among species provides the first attempt to our knowledge to bridge macro- to micro-evolutionary processes in a cetacean group. This perspective across evolutionary time-scales can shed light on the isolation dynamics and their drivers across the speciation continuum of the Phocoenidae.

### 4.1. New insights into the biogeography of the Phocoenidae

The biogeographical history of cetacean species has been hypothesized to results of a succession of vicariant and dispersal events influenced by geological, oceanic, and climatic reorganization during the Late Miocene, Pliocene and early Pleistocene (McGowen et al., 2009; Steeman et al., 2009). Changes in climate, ocean structure, circulation, and marine productivity opened new ecological niches, enhanced individual dispersal and isolation, and fostered specialization to different food resources. All these factors promoted the adaptive radiation in cetaceans which led to the extant species diversity in the odontocete families (Slater et al., 2010). For example, the Monodontidae and Delphinidae are the closest relatives to the Phocenidae. They originated during the Miocene and displayed an accelerated evolution marked with the succession of speciation events during 3 Myr, leading to the extant species diversity in these groups (McGowen et al., 2009; Steeman et al., 2009). The time calibrated phylogeny of the Phocoenidae (Fig. 2) suggests that porpoises also diversified following similar processes during the late Miocene until the early-Pleistocene (between 6 and 2 Myr). This timing is about 2 to 3 Myr more recent than those proposed by McGowen et al. (2009) and Steeman et al. (2009). The increase of the genetic information, node calibrations and number of sequences per species are known to influence phylogenetic inferences and the divergence time estimates (Duchêne et al., 2011; Zheng and Wiens, 2015). The use of complete mitogenomes, two node calibrations (instead of one) and several sequences per species in our study likely explain the difference compare to previous studies.

Consistent with previous findings (Fajardo-Mellor et al., 2006; McGowen et al., 2009; Rosel et al., 1995b), the finless porpoises were the first species to split among the Phocoenidae. As the vast majority of the porpoise fossils found so far come from tropical or subtropical regions (Fajardo-Mellor et al., 2006), and considering their current affinity for warm waters, the finless porpoises seem to be the last members of a group of porpoise species that adapted primarily to tropical waters. Interestingly, finless porpoises further diversified and colonized more temperate waters of the Yellow Sea and Sea of Japan (Fig. 1a). The five remaining porpoise species diverged ∼4.0 Myr ago and all but the vaquita occupy temperate regions with an antitropical distribution (Fig. 1a). Harbor and Dall’s porpoises inhabit the cold water of the Northern hemisphere whereas spectacled and Burmeister’s porpoises are found in the Southern hemisphere. This result is consistent with the hypothesis that antitropically distributed cetaceans have evolved with the deep environmental changes that occurred during the late Pleistocene and as a response to the fluctuations in surface water temperatures in the tropics, concomitant with the changes in oceanographic currents, marine productivity, and feeding areas (Barnes, 1985; Davies, 1963; Pastene et al., 2007). About 3.2 Myr ago, the formation of the Panama Isthmus altered the tropical currents and water temperatures in coastal regions of the Pacific. It promoted the dispersion of numerous taxa from the Northern to the Southern Hemisphere (Lindberg, 1991). This process is also a plausible driver that led to the antitropical distribution of the modern porpoises (Fajardo-Mellor et al., 2006).

During the porpoises’ evolutionary history, a symmetric evolution took place approximatively at the same time in both hemispheres resulting in analogous ecological adaptations. In the Northern hemisphere, the split between Dall’s and harbor porpoises ∼3.1 Myr ago led to offshore *versus* coastal specialized species, respectively. This pattern was mirrored in the Southern hemisphere with the split between the spectacled and Burmeister’s porpoises ∼2.1 Myr ago which led also to the divergence between offshore *versus* coastal specialized species. Such a symmetric habitat specialization likely reflects similar ecological opportunities that opened in both hemispheres and triggered a convergent adaptation in the porpoises. Interestingly, this parallel offshore evolution observed in Dall’s and spectacled porpoises was accompanied by a convergent highly contrasted countershading coloration pattern with a white ventral side and black dorsal back side in both species. This color pattern is thought to be an adaptation to the offshore environment serving as camouflage for prey and predators (Perrin, 2009).

Resources and diet specializations are known to be a major driver of cetacean evolution as their radiation is linked to the colonization of new vacant ecological niches in response to past changes (Slater et al., 2010). As small endothermic predators with elevated energetic needs, limited energy storage capacity and a rapid reproductive cycle, porpoises are known for their strong dependency on food availability (Hoekendijk et al., 2017; Koopman et al., 2002). These characteristics reinforce the hypothesis that their adaptive radiation has been strongly shaped by historical variation in food resources and should also be visible at the intraspecific level.

### 4.2. Porpoises phylogeography and microevolutionary processes

The processes shaping the evolution of porpoises at the macro-evolutionary time scale find their origins at the intraspecific level (micro-evolutionary scale), with the split of multiple lineages within species that may or may not evolve into fully distinct and reproductively isolated species. We showed that all lineages forming the intraspecific subdivisions (sub-species or ecotypes) were all monophyletic groups. All these distinct lineages found their most recent common ancestor within the last million years. These results corroborate previous phylogeographic studies suggesting that intraspecific subdivisions observed in many porpoise species were mediated by environmental changes during the last glacial cycles of the Quaternary (Chen et al., 2017; Fontaine, 2016; Hayano et al., 2003; Zhou et al., 2018). The Last Glacial Maximum (LGM) and the subsequent ice retreat have profoundly shaped the phylogeographic patterns of many organisms, leaving behind multiple divergent lineages in many cetacean taxa that are vestiges of past environmental variations (Chen et al., 2017; Fontaine, 2016; Foote et al., 2016; Louis et al., 2014). Adaptive evolution to different niches in response to past changes associated with variation in marine sea ice, primary production, and redistribution of feeding areas led to intraspecific divergence in many cetaceans in terms of genetics, behavior, morphology, and geographical distribution. For example, the specialization between coastal *versus* offshore ecotypes in bottlenose dolphins has been dated back to the LGM, and explain the observed patterns of genetic and morphological differentiation (Louis et al., 2014). Likewise, the behavior, size, color patterns and genetic divergence among some different types of killer whales were attributed to specialization onto different food resources since the LGM (Foote et al., 2016). The present study shows that analogous processes occurred in each porpoise species too.

The finless porpoises represent probably one of the best documented case of incipient speciation related to habitat specializations among the porpoises. Consistently with the results of Zhou et al. (2018) based on whole genome sequencing, our mitogenome results dated the radiation of the finless porpoise species within the last ∼0.5 Mya, which coincide with profound environmental changes associated with the Quaternary glaciations. In particular, our results are congruent with the hypothesis suggesting that the diversification of the finless porpoises has been driven by the elimination of the Taiwan Strait associated with the sea-level drop during glacial periods (Wang et al., 2008). Indeed, at least three land bridges connected Taiwan to the mainland China since the last 0.5 Myr and could have enhanced the separation between the Indo-Pacific and the narrow ridged finless porpoises (Wang et al., 2008). Likewise, we dated the emergence of the Yangtze finless porpoise to the last ∼0.1 Myr, which is consistent with previous studies suggesting that the last glacial event have strongly determined the evolutionary history of this species (Chen et al., 2017; Zhou et al., 2018).

Similar to the finless porpoises, the harbor porpoises are also divided into several lineages previously recognized as distinct sub-species. The deepest split is observed between the North Pacific (*P. p. vomerina*) and North Atlantic lineages, and is deeper than the genetic divergence observed between the two species of finless porpoises. The lack of shared haplotypes between Pacific and Atlantic porpoises confirm previous results supporting their total isolation (Rosel et al., 1995a). Their splitting time was estimated at ∼0.86 Mya, which is consistent with the presumed time when the North Pacific porpoises colonized the North Atlantic basin (Tolley and Rosel, 2006). The two ocean basins were last in contact across the Arctic approximately 0.7 to 0.9 Myr. ago, with estimated sea surface temperatures of *ca.* 0.5°C (Harris, 2005), which corresponds to the lowest temperature at which harbor porpoises are currently observed. Within the North Atlantic, the three known subspecies (Fontaine, 2016) (i.e. *P. p. phocoena*; *P. p. meridionalis* and *P. p. relicta*) were also detected as distinct monophyletic groups based on the mitogenome (Fig. 1b and S2). Their evolutionary history has been strongly influenced by recent environmental changes during the LGM (reviewed in Fontaine, 2016). North Pacific porpoises also showed unreported cryptic subdivisions (i.e. NP1 and NP2 in Fig. 1b and S2), displaying a level of divergence deeper than the one observed between the Iberian and Mauritanian harbor porpoises or between the Yangtze and East Asian finless porpoises. These two clades (NP1 and NP2) may also represent lineages that split during the LGM. The steady increase in genetic diversity observed in the NP1 lineage since 12 kyrs ago (Fig. 5c) is consistent with the end of the LGM.

Compared to the finless and harbor porpoises, little is known about the Dall’s, Burmeister’s and spectacled porpoises. This is in part due to the limited number of observations and access to biological data (but see Hayano et al., 2003; Pimper et al., 2012; Rosa et al., 2005), especially for the spectacled porpoise. Despite these limitations, our study revealed that the patterns and processes described for the finless porpoise and harbor porpoise may apply also to the majority of the other porpoise species. Previous studies identified multiple intraspecific subdivisions within the Dall’s (Hayano et al., 2003) and Burmeister’s (Rosa et al., 2005) porpoises. The long branches in the phylogenetic tree (Fig. 1b) for Dall’s porpoise (DP1/DP2 lineages) and spectacled porpoises (SP1/SP2) imply that distinct evolutionary units may also exist in these species. Furthermore, their vast distribution (Fig 1a) and divergence times among lineages (Tables S5) suggest that the different lineages in these species also split during the Quaternary glaciations.

The vaquita contrasts strikingly with the other species with its narrow distribution, the smallest of all marine mammals (see Fig. 1a and Lundmark, 2007). Previous studies based on a short fragment of the mitochondrial *D-loop* and *Cyt-b* identified the Burmeister’s porpoise as the closest relative to the vaquitas. However, the phylogenetic results reported here using the whole mitogenome support that the vaquita coalesce with the ancestor of the Burmeister’s and spectacled porpoise with maximal support. The estimated split time of ∼2.4 Myr ago between vaquita and the southern porpoises is consistent with the onset of the Quaternary glaciations, 2.6 Myr ago (Ehlers and Gibbard, 2014). The fact that the vaquita is found in the northern hemisphere, while the Burmeister’s and spectacled porpoises are in the southern hemisphere implies that some ancestors with cold-affinities from the southern species crossed the equator in the past and became isolated long enough to become the ancestors of the extant vaquita. The most parsimonious hypothesis is that the decrease in water temperature associated with a glacial maximum likely allowed vaquita’s ancestor to cross the equator and disperse to the Northern Hemisphere (Norris and McFarland, 1958). The current vaquita representatives form thus a relic population of the temperate southern species’ ancestor that crossed the intertropical zone. In contrast with previous mitochondrial studies that found no variation at the D-loop (Rosel and Rojas-Bracho, 1999) we observed 16 sites segregating along the entire mitogenome. Among them, 13 were singleton mutations. This extremely low nucleotide diversity was the lowest of all porpoise lineages, as illustrated by the extremely short branches in the phylogenetic tree (Fig. 1b and 2). The origin of the present mitochondrial diversity is also relatively recent, with a *T_MRCA_* estimated at ∼20,000 to 70,000 years with the phylogenetic approach and ∼1,500 to 5,000 years with coalescent approach. The main reason behind this discrepancy is called “time dependency of molecular rates” (Ho et al., 2011). Population genetics coalescent-based estimates reflect the spontaneous mutation rates, whereas phylogeny-based estimates reflect the substitution rates (i.e. mutations fixed among taxa). Contrasting with these recent estimates, the branch connecting to the ancestor of vaquitas and southern species dates back to ∼2.4Mya. This suggest that either additional lineages may have existed in the past and went extinct or that only a single lineage crossed the intertropical zone. Whole genome analyses may help enlightening the evolutionary history of this peculiar species.

### 4.3. Genetic diversity and conservation

Maintenance of genetic diversity has been considered as an important factor in conservation biology. Genetic factors can contribute to the “extinction vortex” by a mutual reinforcement with demographic processes speeding-up population decline and increasing their extinction risk (Allendorf et al., 2012). As a consequence, ideal conservation measures should be designed to maximize genetic diversity, especially through the management of evolutionary significant units (ESU) (Moritz, 1994). However, conservation status does not explicitly take this parameter into account since the relationship between IUCN status and genetic diversity is not always straightforward (Nabholz et al., 2008). In this study, the genetic diversity of each porpoise species correlates well with its IUCN status, especially when we account for intraspecific subdivision. The *Critically Endangered* taxa, such as the vaquitas or the YPF porpoises displayed extremely low *π* suggesting a low *Ne*. The *Endangered* (EAF) and *Vulnerable* (Black Sea harbor porpoise) taxa displayed low to average *π*. *Least Concern* taxa (e.g. North Atlantic harbor porpoise and DP2) exhibited higher *π*, suggesting larger *Ne* and/or the presence of internal subdivision. This link between *π* and the IUCN conservation status may thus provide a useful proxy to assess the conservation status of taxa for which an IUCN status has not been yet established, due to data deficiency. For example, the Iberian harbor porpoise population is among the marine mammals displaying the highest stranding and by-catch rates reported (Read et al., 2012). The low genetic diversity reported in the present study represents thus an additional signal indicating how possibly vulnerable are these Iberian porpoises. On the other hand, spectacled porpoises represent currently one of the least known cetacean species. Their genetic diversity is comparable to the one observed in the Dall’s porpoise DP2 lineage or the North Atlantic (NAT) harbor porpoise linage, suggesting these populations display large *Ne*.

Mitochondrial diversity may not always be a good proxy of population abundance. Other evolutionary processes than just demography may impact genetic diversity (i.e., such as natural selection) (Bazin et al., 2006). The *d_N_*/*d_S_* and *π_N_/π_S_* lower than 1 highlighted in this study would be usually indicative of purifying selection acting on the mitochondrial genetic diversity. The negative relationships observed between *π* and *d_N_*/*d_S_* (Fig. S4b) lend further support to this hypothesis, suggesting that purifying selection is more effective in large populations as predicted by the neutral theory (Hughes, 2008; Phifer-Rixey et al., 2012). Surprisingly, the MK-tests suggested that purifying selection was prevailing on the mitochondrial genomes of the endangered porpoises with *NI* values larger than 1. In contrast, selective neutrality could not be rejected for less threatened species. At first glance, this result for the endangered porpoises seems counter intuitive. Purifying selection is expected to be less effective in lineages where population size is very small, since genetic drift is expected to outperform selective forces (Kimura, 1983). However, when a lineage harbors a low *Ne*, slightly deleterious variants are expected to increase in frequency and segregate for a longer time without being fixed. Parsch et al. (2009) showed that this effect will increase the number of polymorphic nonsynonymous mutations compared to the divergent nonsynonymous mutations (*π_N_ >> d_N_*). Hence, despite the reported increase of *d_N_*/*d_S_* in endangered taxa, it induces a bias in the neutrality index toward positive values creating an artefactual signal of purifying selection. The higher values of *π_N_/π_S_* observed in the endangered porpoise taxa (Table S3) and the negative relationships reported between *π* and *NI* support this hypothesis. All these elements suggest that demographic processes rather than selective forces drive the genetic diversity of the mitochondrial genome, and led to high values of NI or *d_N_*/*d_S_* in the endangered taxa.

## 5. Conclusions

Using complete mitochondrial genomes, we reconstructed the most comprehensive picture of the evolutionary history of the Phocoenidae to date. Besides clarifying the debated phylogenetic relationships among porpoises, our results provided new insights into the process driving species diversification in the porpoises across the speciation continuum. Similar to the Delphinidae, the Phocoenidae recently radiated in response to past environmental variation, adapting to different environments, ecological niches, and food resources. Furthermore, our results suggested that the processes governing their divergence at the macro-evolutionary scale find their origins at the micro-evolutionary scale. We revealed unreported genetic subdivision for several taxa suggesting that our knowledge about many species, especially the data deficient southern species, is still scarce. Finally, the level of mitochondrial genetic diversity within a species seems to be primarily driven by demographic processes, rather than natural selection and turned out to be a good proxy for the conservation issues reported in these groups (i.e. Yangtze finless porpoises or vaquita). In the future, the analyses of genome-wide data and a better description of porpoise distinctive environments hold great promise to shed light on the processes underpinning their adaptation and divergence. Such knowledge can facilitate the identification of units following distinct evolutionary trajectories, which in turn can greatly improve our ability to set conservation plans to mitigate the decline of several porpoise groups.

## Acknowledgments

The following persons deserve our special thanks for having collected and made the samples available for this study: Tom Jefferson, Carlos Olavarria, Jay Barlow, Robin Baird, Marilyn Dahlheim, Lorenzo Rojas-Bracho. We thank Andrew D. Foote for his constructive feedbacks that greatly improved the manuscript. We are also grateful to Frédéric Labbé for his assistance with the preparation of the map in Fig. 1 and Jorge Eduardo Amaya Romero for helping with the MITOBIM pipeline. We would like also to thank the Center for Information Technology of the University of Groningen for their support and for providing access to the *Peregrine* high-performance computing cluster. This study benefitted from the funding of the University of Groningen (The Netherlands). YBC was supported by a PhD fellowship from the University of Groningen. MF was supported through a Portuguese Foundation for Science and Technology (FCT) grant SFRH/BD/30240/2006.

## Author’s contributions

MCF designed the study with the contribution of PAM; YBC analyzed the data under the supervision of MCF, with help from AW and JR; AA, AB, MF, BLT, LRB, KMR, GAV, PAM contributed with the biological materials; JT, KMR performed the DNA extractions; CS and TH constructed the genomic libraries at Swift Bioscience and performed part of the sequencing; YBC and MCF wrote the manuscript with input and feedbacks from all the co-authors.

## Data accessibility

Mitochondrial genomes were deposited on NCBI under the accession numbers To Be Announced (TBA). Data and scripts have also been deposited on Dryad doi: TBA

