## Supplementary Materials for "Evolutionary history of the porpoises (*Phocoenidae*) across the speciation continuum: a mitogenome phylogeographic perspective"

### Index

### Supplementary text

**Text S1. Reads quality check and filtering.**

An initial quality check (QC) of the reads was conducted using *FastQC* v.0.11.5. Specifically, we assessed the general phred-scale quality score of the reads (on average and for every site of the reads), the presence of overrepresented sequences or of Illumina specific adaptors in the raw data. General quality statistics of the raw data are provided in Table S2. Then, low-quality reads, overrepresented sequences and Illumina adaptors were removed using *Trimmomatic* v0.36 (Bolger et al., 2014). We applied two different sets of filters according to the two distinct sequencing technology used (MySeq or HiSeq). For the 44 Miseq sequences and the finless porpoise short read archive (Yim et al., 2014) from NCBI, we used the following command in *Trimmomatic*: [ILLUMINACLIP:/TruSeq3-PE.fa:2:30:10 LEADING:5 TRAILING:5 SLIDINGWINDOW:4:15 MINLEN:50 HEADCROP:4]. This performed the following operations:

1. Remove the Illumina adaptors using the "IlluminaClip" option.
2. Remove low phred-scale quality score bases (Q < 5) at the beginning (LEADING option) and ending (TRAILING option) of each read.
3. Slide a 4 base-pairs (bps) window along the reads and replace by N's if the average quality lower or equal to 15.
4. Require a minimal read length of 50 bp (before using the HEADCROP option of *Trimmomatic* v0.36 in 5.) otherwise we dropped it.
5. Remove the first 4 bases (to improve the "Sequence content" statistics of *FastQC*).

For the reads generated using a Hiseq-4000 (at BGI) for the 12 harbor porpoises, we used the following command in *Trimmomatic*: [ILLUMINACLIP:/TruSeq3-PE.fa:2:30:10 LEADING:20 TRAILING:20 SLIDINGWINDOW:4:15 MINLEN:50 HEADCROP:4]. This performed the following operations:

1. Remove the Illumina adaptors using the "IlluminaClip" option.
2. Remove low phred-scale quality score bases (inferior to 20) at the beginning (LEADING option) and at the ending (TRAILING option) of each read.
3. Slide a 4 bps window along the reads and replace by N's if the average quality lower or equal to 15.
4. Require a minimal read length of 50 bp (to improve the "Sequence content" statistics of *FastQC*.), otherwise we dropped it.
5. Remove the first 4 bases (to improve the "Sequence content" statistics of *FastQC*).

A post-cleaning QC assessment showed that the average quality score of the data increased dramatically (Table S2) and no adaptor, nor duplicated reads were detected after the data cleaning step. The number of reads kept and other general statistics about the data after QC filtering are provided in Table S2.

**Text S2. Mitogenome assembly using MITOBIM.**

As a second approach to assemble mitogenomes from short reads, we used the baiting and iterative elongation procedure of MITOBIM (Hahn et al., 2013). This method involves two steps. First, the reads are mapped to a reference genome using *MIRA* v4.0 (Chevreux et al., 1999), generating a new reference genome based only on the most conserved regions. Secondly, the remaining reads that overlap with the new reference genome are iteratively fished and mapped to the reference genome using *MITOBIM* v1.8 (Hahn et al., 2013). At each iteration, this process extends the new reference until reaching a stationary number of reads. The first step has significant memory requirements and leads to a significant increase of the computational time. This increased memory consumption can be bypassed using the quick option of *MITOBIM* by skipping the first steps that uses *MIRA* (Hahn et al., 2013). This option reduces the total pool of reads only to the ones that have certain k-mers overlap (≥ 31 bp) to the reference already before the initial assembly. This approach performs well with samples that are not too distantly related from the references available (See details in Hahn et al., 2013). We assembled two individuals with the "classical approach" (*MIRA* and *MITOBIM*) and with the quick option to compare the results of both methods. As the results were identical, we assembled the 55 remaining samples using only the quick option. The statistics of the assembly with *MITOBIM* is provided in Table S2.

We compared visually the assemblies generated by *Geneious* and MITOBIM in *Geneious*. When the assembly with *Geneious* led to ambiguous sites (IUPAC ambiguity codes), we took the nucleotides called by *MITOBIM*. When disagreements were observed between assemblies, we checked the coverage and visually inspected the reads mapping at those sites in *Geneious*. If the coverage was less than 5, we used the base attributed by *MITOBIM*. If the coverage was more than 5 and the majority (at least 75%) of the reads mapped at that site (in *Geneious*) contained a certain base, we used it.

### Supplementary tables

#### Table S1. Sampling information by individual.

| Species | Source[1] | ID | Sex[2] | Year | Latitude | Longitude | Locality |
|---|---|---|---|---|---|---|---|
| *N. pho.* | SWFSC | 983 | U | NA | NA | NA | China, Yangtze River |
| *N. pho.* | SWFSC | 984 | NA | NA | NA | NA | NA |
| *N. pho.* | SWFSC | 7859 | NA | NA | NA | NA | NA |
| *N. pho.* | SWFSC | 7869 | NA | NA | NA | NA | NA |
| *N. pho* | SWFSC | 9559 | F | 1996 | 22.74 | 120.21 | Taiwan, Yung-An |
| *N. pho* | SWFSC | 9560 | NA | NA | NA | NA | NA |
| *P. dal.* | SWFSC | 145217 | U | 2014 | 47.87 | -125.15 | NA |
| *P. dal.* | SWFSC | 145412 | U | 2014 | 33.35 | -120.33 | NA |
| *P. dio.* | SWFSC | 981 | U | 1989 | -53.1 | -66.5 | Argentina, Estancia Las Violetas |
| *P. dio.* | SWFSC | 7014 | F | 1997 | -42.9 | 148.4 | Australia, Tasmania, Bruny Is., Adventure Bay |
| *P. dio.* | SWFSC | 7015 | NA | NA | NA | NA | NA |
| *P. pho.* | SWFSC | 161 | M | NA | 36.80 | -121.78 | USA, Ca, Moss Landing |
| *P. pho.* | SWFSC | 704 | F | 1989 | 36.90 | -121.83 | USA, Ca, Monterey Bay |
| *P. pho.* | SWFSC | 706 | F | 1989 | 36.80 | -121.78 | USA, Ca, Monterey |
| *P. pho.* | SWFSC | 707 | M | 1989 | 36.90 | -121.83 | USA, Ca, Santa Cruz Co., Sunset State Beach |
| *P. pho.* | SWFSC | 1080 | F | 1988 | 49.05 | -125.72 | Canada, British Columbia, Long Beach |
| *P. pho.* | SWFSC | 1082 | M | 1990 | 48.42 | -123.40 | Canada, British Columbia, Victoria |
| *P. pho.* | SWFSC | 1084 | F | 1987 | 49.00 | -123.50 | Canada, British Columbia, Gabriola Is. |
| *P. pho.* | SWFSC | 3766 | M | 1989 | 36.63 | -121.87 | USA, Ca, (Northern) |
| *P. pho.* | SWFSC | 3767 | M | 1989 | 36.55 | -121.98 | USA, Ca, (Northern) |
| *P. pho.* | SWFSC | 3768 | M | 1989 | 36.63 | -121.87 | USA, Ca, (Northern) |
| *P. pho.* | SWFSC | 5411 | F | 1993 | 57.03 | -154.15 | USA, Ak, Kodiak |
| *P. pho.* | SWFSC | 8514 | F | 1994 | 42.83 | -124.58 | USA, Or, Port Orford |

| | | | | | | | |
|---|---|---|---|---|---|---|---|
| *P. pho.* | SWFSC | 4764 | NA | NA | NA | NA | NA |
| *P. pho.* | SWFSC | 26609 | NA | NA | NA | NA | NA |
| *P. pho.* | Fontaine *et al.* 2014 | 2000-24 | M | 2000 | 66.5 | 12.08 | Norway |
| *P. pho.* | Fontaine *et al.* 2014 | IFR4 | M | 1990 | -6.90 | 62.20 | Faroe Islands |
| *P. pho.* | Fontaine *et al.* 2014 | PP118-2004 | M | 2004 | 40.15 | -8.8667 | Portugal |
| *P. pho.* | Fontaine *et al.* 2014 | PP30-2002 | M | 2002 | 40.3167 | -8.86 | Portugal |
| *P. pho.* | Fontaine *et al.* 2014 | PP79-2003 | M | 2003 | 40.4333 | -8.86 | Portugal |
| *P. pho.* | Fontaine *et al.* 2014 | RIM100 | U | NA | 19 | -17 | Mauritania |
| *P. pho.* | Fontaine *et al.* 2014 | RIM156 | M | NA | 19 | -17 | Mauritania |
| *P. pho.* | Fontaine *et al.* 2014 | RIM99 | M | NA | 19 | -17 | Mauritania |
| *P. pho.* | Fontaine *et al.* 2014 | SV276 | M | NA | -21.95 | 64.27 | Iceland |
| *P. pho.* | Fontaine *et al.* 2014 | U64 | M | 1998 | 44.6 | 33.53 | Ukraine, Black Sea |
| *P. pho.* | Fontaine *et al.* 2014 | U75 | F | 1998 | 44.6 | 33.53 | Ukraine, Black Sea |
| *P. pho.* | Fontaine *et al.* 2014 | U93 | M | 1998 | 44.6 | 33.53 | Ukraine, Black Sea |
| ***P. sin.*** | SWFSC | 703 | U | 1985 | 34.42 | -120.50 | Mexico, El Burro |
| *P. sin.* | SWFSC | 1649 | U | 1993 | 31.75 | -114.50 | Mexico, Gulf Of California |
| *P. sin.* | SWFSC | 1651 | U | 1993 | 31.75 | -114.50 | Mexico, Gulf Of California |
| *P. sin.* | SWFSC | 1654 | F | 1992 | 31.75 | -114.50 | Mexico, Baja California North, El Ouelele |
| *P. sin.* | SWFSC | 1655 | M | 1993 | 31.75 | -114.58 | Mexico, Sonora, Gulf Of Santa Clara |
| *P. sin.* | SWFSC | 1660 | F | 1993 | 31.75 | -114.58 | Mexico, Sonora, Gulf Of Santa Clara |
| *P. sin.* | SWFSC | 4018 | U | 1980 | 31.75 | -114.50 | Mexico, Upper Gulf Of Ca |
| *P. sin.* | SWFSC | 4379 | F | 1990 | 31.75 | -114.50 | Mexico, Gulf Of Santa Clara |
| *P. sin.* | SWFSC | 4381 | M | 1990 | 31.75 | -114.50 | Mexico, Gulf Of Santa Clara |
| *P. sin.* | SWFSC | 4393 | F | 1991 | 31.75 | -114.50 | Mexico, Gulf Of Santa Clara |
| *P. sin.* | SWFSC | 4394 | M | 1991 | 31.75 | -114.50 | Mexico, Gulf Of Santa Clara |
| *P. sin.* | SWFSC | 4396 | F | 1991 | 31.75 | -114.50 | Mexico, Gulf Of Santa Clara |
| *P. spi.* | SWFSC | 1092 | U | 1990 | -15 | -76 | Peru |

| | | | | | | | |
|---|---|---|---|---|---|---|---|
| *P. spi.* | SWFSC | 52776 | F | 1997 | -32.92 | -71.52 | Chile, Mantagua |
| *P. spi.* | SWFSC | 52777 | M | 1999 | -35.33 | -72.42 | Chile, Constitucion |

NA, Missing information ; [1]SWFSC, Southwest Fisheries Science Centre, NOAA; Fontaine et al. (2014) ; [2]F: Female; M: Male; U: Unknown.

#### Table S2. Reads filtering and mitochondrial assembly using *MITObim* and *Geneious*.

| General information | | | | | Reads before cleaning | | | Reads after cleaning | | | | | MITOBIM | | | | |
|---|---|---|---|---|---|---|---|---|---|---|---|---|---|---|---|---|---|
| **ID** | **Sequencing techology** | **Species** | **Data generation** | **GenBank** | **Reads** | **Reads length** | **%GC** | **Reads** | **%GC** | **Reads length** | **Average read length F.** | **Average read length R.** | **Assembly size** | **Reads mapped** | **Average coverage** | **%GC in assembly** | **Mapped reads** |
| 1092 | Miseq | *P. spinipinnis* | This study | TBA | 2307042 | 101 | 38 | 2262214 | 38 | 46-97 | 95.81 | 95.59 | 16402 | 25260 | 157.90 | 40 | 1.1 |
| 52776 | Miseq | *P. spinipinnis* | This study | TBA | 2069602 | 101 | 40 | 2037416 | 40 | 46-97 | 95.78 | 95.31 | 16390 | 3429 | 24.05 | 40 | 0.1 |
| 52777 | Miseq | *P. spinipinnis* | This study | TBA | 2672488 | 101 | 40 | 2628532 | 40 | 46-97 | 95.75 | 95.02 | 16392 | 7132 | 46.72 | 40 | 0.2 |
| 145216 | Miseq | *P. dalli* | This study | TBA | 3214672 | 101 | 41 | 3162846 | 41 | 46-97 | 95.74 | 95.03 | 16383 | 4758 | 32.16 | 41 | 0.1 |
| 145217 | Miseq | *P. dalli* | This study | TBA | 2279936 | 101 | 41 | 2235812 | 41 | 46-97 | 95.77 | 95.22 | 16388 | 4319 | 29.51 | 41 | 0.1 |
| 145242 | Miseq | *P. dalli* | This study | TBA | 4510470 | 101 | 41 | 4439616 | 41 | 46-97 | 95.74 | 95.05 | 16387 | 6960 | 45.69 | 41 | 0.1 |
| 145258 | Miseq | *P. dalli* | This study | TBA | 2124314 | 101 | 41 | 2078282 | 41 | 46-97 | 95.73 | 94.71 | 16367 | 3415 | 23.92 | 41 | 0.1 |
| 145412 | Miseq | *P. dalli* | This study | TBA | 2566400 | 101 | 41 | 2522672 | 41 | 46-97 | 95.76 | 95.23 | 16367 | 4511 | 30.74 | 41 | 0.1 |
| 145413 | Miseq | *P. dalli* | This study | TBA | 2067734 | 101 | 41 | 2037046 | 41 | 46-97 | 95.74 | 95.12 | 16381 | 2733 | 19.76 | 41 | 0.1 |
| 7859 | Miseq | *N. phocaenoides* | This study | TBA | 2347308 | 101 | 43 | 2298100 | 42 | 46-97 | 95.69 | 95.09 | 16385 | 4703 | 31.87 | 41 | 0.2 |
| 7869 | Miseq | *N. phocaenoides* | This study | TBA | 2563718 | 101 | 42 | 2523568 | 42 | 46-97 | 95.71 | 95.12 | 16385 | 4231 | 28.96 | 41 | 0.1 |
| 9559 | Miseq | *N. phocaenoides* | This study | TBA | 1840894 | 101 | 40 | 1809462 | 40 | 46-97 | 95.79 | 95.35 | 16385 | 6551 | 43.26 | 41 | 0.3 |
| 9560 | Miseq | *N. phocaenoides* | This study | TBA | 2269316 | 101 | 40 | 2235780 | 40 | 46-97 | 95.77 | 95.14 | 16385 | 7981 | 51.89 | 41 | 0.3 |
| 983 | Miseq | *N. phocaenoides* | This study | TBA | 2082960 | 101 | 41 | 2049768 | 40 | 46-97 | 95.77 | 95.30 | 16386 | 26081 | 162.93 | 41 | 1.2 |
| 984 | Miseq | *N. phocaenoides* | This study | TBA | 2467902 | 101 | 40 | 2421426 | 40 | 46-97 | 95.76 | 95.12 | 16386 | 11559 | 73.91 | 41 | 0.4 |
| 1080 | Miseq | *P. phocoena* | This study | TBA | 5095324 | 101 | 41 | 4997006 | 41 | 46-97 | 95.70 | 93.87 | 16384 | 7611 | 49.29 | 41 | 0.1 |
| 1082 | Miseq | *P. phocoena* | This study | TBA | 5628180 | 101 | 41 | 5506628 | 41 | 46-97 | 95.69 | 93.90 | 16384 | 2423 | 17.74 | 41 | 0.0 |
| 1084 | Miseq | *P. phocoena* | This study | TBA | 6099638 | 101 | 41 | 5990164 | 41 | 46-97 | 94.10 | 95.48 | 16384 | 4311 | 29.20 | 41 | 0.0 |
| 161 | Miseq | *P. phocoena* | This study | TBA | 6162988 | 101 | 42 | 6043918 | 41 | 46-97 | 95.68 | 93.98 | 16384 | 3018 | 21.40 | 41 | 0.0 |
| 26609 | Miseq | *P. phocoena* | This study | TBA | 2320654 | 101 | 41 | 2279300 | 41 | 46-97 | 95.75 | 95.15 | 16385 | 3147 | 22.28 | 41 | 0.1 |
| 3766 | Miseq | *P. phocoena* | This study | TBA | 7919712 | 101 | 42 | 7388012 | 41 | 46-97 | 95.65 | 93.84 | 16384 | 33018 | 204.21 | 41 | 0.4 |
| 3767 | Miseq | *P. phocoena* | This study | TBA | 7444496 | 101 | 42 | 6907630 | 42 | 46-97 | 95.64 | 94.13 | 16384 | 26015 | 161.86 | 41 | 0.3 |
| 3768 | Miseq | *P. phocoena* | This study | TBA | 7230284 | 101 | 42 | 7009836 | 42 | 46-97 | 95.66 | 94.06 | 16384 | 17549 | 110.10 | 41 | 0.2 |
| 4764 | Miseq | *P. phocoena* | This study | TBA | 2863216 | 101 | 41 | 2816740 | 40 | 46-97 | 95.76 | 95.06 | 16385 | 4288 | 29.25 | 41 | 0.1 |
| 5411 | Miseq | *P. phocoena* | This study | TBA | 3649262 | 101 | 41 | 3549554 | 42 | 46-97 | 95.69 | 93.39 | 16385 | 6985 | 45.51 | 41 | 0.2 |
| 704 | Miseq | *P. phocoena* | This study | TBA | 5656192 | 101 | 41 | 5535084 | 41 | 46-97 | 95.71 | 93.69 | 16384 | 4342 | 29.41 | 41 | 0.0 |
| 706 | Miseq | *P. phocoena* | This study | TBA | 4224202 | 101 | 41 | 4127958 | 41 | 46-97 | 95.68 | 93.78 | 16384 | 3187 | 22.34 | 41 | 0.0 |
| 707 | Miseq | *P. phocoena* | This study | TBA | 5746290 | 101 | 42 | 5639666 | 41 | 46-97 | 95.68 | 93.89 | 16384 | 7411 | 48.12 | 41 | 0.1 |
| 8514 | Miseq | *P. phocoena* | This study | TBA | 4618658 | 101 | 41 | 4533816 | 41 | 46-97 | 95.69 | 93.90 | 16384 | 5839 | 38.60 | 41 | 0.1 |
| 7014 | Miseq | *P. dioptrica* | This study | TBA | 2159390 | 101 | 41 | 2126000 | 40 | 46-97 | 95.78 | 95.30 | 16371 | 4247 | 29.10 | 40 | 0.2 |
| 7015 | Miseq | *P. dioptrica* | This study | TBA | 2451054 | 101 | 40 | 2411570 | 40 | 46-97 | 95.72 | 95.09 | 16383 | 2431 | 17.91 | 40 | 0.1 |
| 981 | Miseq | *P. dioptrica* | This study | TBA | 2518274 | 101 | 40 | 2463666 | 40 | 46-97 | 95.78 | 95.41 | 16371 | 9630 | 62.33 | 40 | 0.3 |
| 1649 | Miseq | *P. sinus* | This study | TBA | 2.1E+07 | 101 | 40 | 20501376 | 40 | 46-97 | 95.70 | 95.16 | 16411 | 46375 | 285.92 | 40 | 0.2 |
| 1651 | Miseq | *P. sinus* | This study | TBA | 1.1E+07 | 101 | 41 | 10677908 | 41 | 46-97 | 95.68 | 95.09 | 16390 | 25302 | 157.80 | 40 | 0.2 |
| 1654 | Miseq | *P. sinus* | This study | TBA | 2296982 | 101 | 41 | 2258892 | 41 | 46-97 | 95.77 | 95.40 | 16388 | 5066 | 34.13 | 40 | 0.2 |
| 1655 | Miseq | *P. sinus* | This study | TBA | 1753050 | 101 | 41 | 1726422 | 41 | 46-97 | 95.77 | 95.32 | 16369 | 3349 | 23.59 | 40 | 0.1 |
| 1660 | Miseq | *P. sinus* | This study | TBA | 2284180 | 101 | 41 | 2252156 | 41 | 46-97 | 95.76 | 95.34 | 16388 | 5387 | 36.09 | 40 | 0.2 |
| 4018 | Miseq | *P. sinus* | This study | TBA | 3306892 | 101 | 40 | 3220874 | 40 | 46-97 | 95.77 | 95.46 | 16389 | 9968 | 64.16 | 40 | 0.3 |
| 4379 | Miseq | *P. sinus* | This study | TBA | 2931966 | 101 | 41 | 2872968 | 40 | 46-97 | 95.77 | 95.37 | 16389 | 7344 | 48.09 | 40 | 0.2 |
| 4381 | Miseq | *P. sinus* | This study | TBA | 4983672 | 101 | 41 | 4880416 | 40 | 46-97 | 95.76 | 95.35 | 16370 | 6004 | 39.97 | 40 | 0.1 |
| 4393 | Miseq | *P. sinus* | This study | TBA | 3801750 | 101 | 40 | 3727950 | 40 | 46-97 | 95.77 | 95.43 | 16387 | 6918 | 45.48 | 40 | 0.1 |
| 4394 | Miseq | *P. sinus* | This study | TBA | 3577624 | 101 | 41 | 3505548 | 41 | 46-97 | 95.73 | 95.13 | 16385 | 5768 | 38.46 | 40 | 0.1 |
| 4396 | Miseq | *P. sinus* | This study | TBA | 6489926 | 101 | 41 | 6348948 | 41 | 46-97 | 95.75 | 95.49 | 16370 | 9066 | 58.76 | 40 | 0.1 |
| 703 | Miseq | *P. sinus* | This study | TBA | 2644732 | 101 | 40 | 2638913 | 40 | 46-97 | 95.76 | NA | 16370 | 2627 | 19.16 | 40 | 0.1 |
| 2000-24 | Hiseq | *P. phocoena* | This study | TBA | 5.6E+07 | 150 | 44 | 51726410 | 44 | 46-146 | 143.46 | 139.73 | 16383 | 89121 | 777.27 | 41 | 0.1 |
| IFR4 | Hiseq | *P. phocoena* | This study | TBA | 5.4E+07 | 150 | 46 | 49604876 | 46 | 46-146 | 143.48 | 139.64 | 16383 | 162993 | 1419.05 | 41 | 0.3 |
| PP118-2004 | Hiseq | *P. phocoena* | This study | TBA | 6.6E+07 | 150 | 44 | 61645074 | 44 | 46-146 | 142.49 | 139.95 | 16384 | 250855 | 2174.67 | 41 | 0.4 |
| PP30-2002 | Hiseq | *P. phocoena* | This study | TBA | 7.3E+07 | 150 | 44 | 67442586 | 43 | 46-146 | 143.35 | 139.84 | 16384 | 364434 | 3167.24 | 41 | 0.5 |
| PP79-2003 | Hiseq | *P. phocoena* | This study | TBA | 5.7E+07 | 150 | 44 | 52438778 | 44 | 46-146 | 143.46 | 139.70 | 16384 | 94088 | 819.13 | 41 | 0.1 |
| RIM100 | Hiseq | *P. phocoena* | This study | TBA | 5.6E+07 | 150 | 45 | 51038992 | 45 | 46-146 | 143.56 | 139.44 | 16385 | 62170 | 543.09 | 41 | 0.1 |
| RIM156 | Hiseq | *P. phocoena* | This study | TBA | 5.8E+07 | 150 | 44 | 52969152 | 44 | 46-146 | 143.51 | 139.70 | 16384 | 77763 | 678.40 | 41 | 0.1 |
| RIM99 | Hiseq | *P. phocoena* | This study | TBA | 6.6E+07 | 150 | 44 | 60786448 | 44 | 46-146 | 143.50 | 139.80 | 16384 | 163977 | 1429.67 | 41 | 0.2 |
| SV276 | Hiseq | *P. phocoena* | This study | TBA | 6.1E+07 | 150 | 45 | 55400950 | 45 | 46-146 | 143.48 | 139.34 | 16384 | 52273 | 456.33 | 41 | 0.0 |
| U64 | Hiseq | *P. phocoena* | This study | TBA | 6.2E+07 | 150 | 44 | 56037790 | 44 | 46-146 | 143.51 | 139.09 | 16382 | 502153 | 4359.84 | 41 | 0.9 |
| U75 | Hiseq | *P. phocoena* | This study | TBA | 6E+07 | 150 | 45 | 54986682 | 45 | 46-146 | 143.48 | 139.44 | 16382 | 274556 | 2388.61 | 41 | 0.5 |
| U93 | Hiseq | *P. phocoena* | This study | TBA | 5.8E+07 | 150 | 44 | 53371864 | 44 | 46-146 | 143.48 | 139.88 | 16382 | 311327 | 2711.72 | 41 | 0.5 |
| SRR940959 | Hiseq | *N. phocaenoides* | Yim *et al.* 2014 | SRR940959 | 5214672 | 100 | 44 | 4992506 | 40 | 46-96 | 94.39 | 94.39 | 16385 | 12425 | 74 | 41 | 0.4 |
| NC_021461 | - | *N. phocaenoides* | Unpublished | NC_021461 | - | - | - | - | - | - | - | - | - | - | - | - | - |
| KR_108307 | - | *N. phocaenoides* | Unpublished | KR_108307 | - | - | - | - | - | - | - | - | - | - | - | - | - |
| KR_108308 | - | *N. phocaenoides* | Unpublished | KR_108308 | - | - | - | - | - | - | - | - | - | - | - | - | - |
| KU886000 | - | *N. phocaenoides* | Unpublished | KU886000 | - | - | - | - | - | - | - | - | - | - | - | - | - |
| KT_852939 | - | *N. phocaenoides* | Cheng *et al.* 2016 | KT_852939 | - | - | - | - | - | - | - | - | - | - | - | - | - |
| AJ55406 | - | *P. phocoena* | Arnason *et al.* 2004 | AJ55406 | - | - | - | - | - | - | - | - | - | - | - | - | - |
| NC_005279 | - | *M. monoceros* | Arnason *et al.* 2004 | NC_005279 | - | - | - | - | - | - | - | - | - | - | - | - | - |
| KF570384 | - | *T. truncatus* | Moura *et al.* 2013 | KF570384 | - | - | - | - | - | - | - | - | - | - | - | - | - |
| KF570385 | - | *T. truncatus* | Moura *et al.* 2013 | KF570385 | - | - | - | - | - | - | - | - | - | - | - | - | - |
| KF570389 | - | *T. truncatus* | Moura *et al.* 2013 | KF570389 | - | - | - | - | - | - | - | - | - | - | - | - | - |
| GU187216 | - | *O. orca* | Morin *et al.* 2010 | GU187216 | - | - | - | - | - | - | - | - | - | - | - | - | - |
| GU187217 | - | *O. orca* | Morin *et al.* 2010 | GU187217 | - | - | - | - | - | - | - | - | - | - | - | - | - |
| GU187218 | - | *O. orca* | Morin *et al.* 2010 | GU187218 | - | - | - | - | - | - | - | - | - | - | - | - | - |
| NC_022805 | - | *T. australis* | Moura *et al.* 2013 | NC_022805 | - | - | - | - | - | - | - | - | - | - | - | - | - |

GenBank: GenBank Accession numbers; Reads: total number of reads, TBA: To be announced

**Table S3. Genetic diversity in the coding regions (13 CDS) of the mitochondrial genome (11,334 bp).**

| | | *N* | *MD* | *H* | $H_d$ (%) | *S* | *Shared P.* | *Singl.* | $\pi$ (%) | $\theta_W$ (%) | *#Syn* | *#NSyn* | $\pi_S$ (%) | $\pi_{NS}$ (%) | $\pi_{NS}/\pi_S$ |
|---|---|---|---|---|---|---|---|---|---|---|---|---|---|---|---|
| **Species** | **All** | 63 | 1 | 56 | 99.4 | 2463 | 2283 | 180 | 6.45 | 4.61 | 2059 | 373 | 21.83 | 1.38 | 0.06 |
| | **FP** | 12 | 1 | 12 | 100.0 | 193 | 46 | 147 | 0.42 | 0.56 | 151 | 42 | 1.28 | 0.13 | 0.10 |
| | **BP** | 3 | 0 | 3 | 100.0 | 25 | 0 | 25 | 0.14 | 0.14 | 21 | 4 | 0.5 | 0.03 | 0.06 |
| | **V** | 12 | 0 | 7 | 86.4 | 10 | 2 | 8 | 0.02 | 0.03 | 8 | 2 | 0.07 | 0.004 | 0.06 |
| | **SP** | 3 | 0 | 3 | 100.0 | 120 | 0 | 120 | 0.70 | 0.70 | 106 | 14 | 2.52 | 0.11 | 0.04 |
| | **DP** | 6 | 0 | 5 | 93.3 | 173 | 49 | 124 | 0.60 | 0.67 | 155 | 18 | 2.13 | 0.09 | 0.04 |
| | **HP** | 27 | 1 | 26 | 99.7 | 474 | 357 | 117 | 1.34 | 1.11 | 384 | 90 | 4.36 | 0.31 | 0.07 |
| **HP** | **NAT** | 4 | 1 | 4 | 100.0 | 82 | 15 | 67 | 0.38 | 0.40 | 66 | 16 | 1.25 | 0.10 | 0.08 |
| | **IB** | 3 | 1 | 3 | 100.0 | 23 | 0 | 23 | 0.14 | 0.14 | 15 | 8 | 0.35 | 0.07 | 0.20 |
| | **MA** | 3 | 1 | 3 | 100.0 | 9 | 0 | 9 | 0.05 | 0.05 | 6 | 3 | 0.14 | 0.03 | 0.21 |
| | **BS** | 3 | 1 | 3 | 100.0 | 14 | 0 | 14 | 0.08 | 0.08 | 8 | 6 | 0.19 | 0.05 | 0.26 |
| | **NP** | 14 | 1 | 13 | 98.9 | 91 | 52 | 39 | 0.23 | 0.25 | 77 | 14 | 0.79 | 0.05 | 0.06 |
| **NP** | **NP1** | 10 | 1 | 10 | 100.0 | 57 | 19 | 38 | 0.14 | 0.18 | 48 | 9 | 0.46 | 0.03 | 0.07 |
| | **NP2** | 4 | 1 | 3 | 83.3 | 3 | 1 | 2 | 0.02 | 0.01 | 3 | 0 | 0.06 | 0 | 0 |
| **FP** | **YFP** | 6 | 1 | 6 | 100.0 | 26 | 3 | 23 | 0.08 | 0.10 | 17 | 9 | 0.22 | 0.04 | 0.17 |
| | **EAF** | 5 | 1 | 5 | 100.0 | 46 | 2 | 44 | 0.17 | 0.20 | 33 | 13 | 0.48 | 0.07 | 0.14 |
| **DP** | **DP2** | 5 | 1 | 4 | 90.0 | 90 | 40 | 50 | 0.38 | 0.38 | 77 | 13 | 1.34 | 0.07 | 0.05 |
| **SP** | **SP2** | 2 | 1 | 2 | 100.0 | 30 | 2 | 30 | 0.27 | 0.27 | 25 | 5 | 0.89 | 0.06 | 0.06 |

FP, *Neophocaena*; BP, *P. spinipninis*, V, *P. sinus*; SP, *P. dioptrica*; DP, *P. dalli*; HP, *P. phocoena*; NAT, North Atlantic harbor porpoise; IB, Iberian harbor porpoise; MA, Mauritanian harbor porpoise; BS, Black Sea harbor porpoise; NP; Pacific *P. phocoena*; YF, *N. a. asiaeorientalis*; EAF, *N. a. sunameri*. For NP1, NP2, DP2 and SP2 see main text Fig. 1b. *N*, Sample size; *MD*, Number of sites with missing data; *H*, number of haplotypes; $H_d$, haplotypic diversity; *S*, segregating sites; *Shared P.*, shared polymorphism; *Singl.*, singleton; $\pi$, average nucleotide diversity by site; $\theta_W$, theta from *S*; *#Syn.*, number of synonymous mutations; *#NSyn.*, number of non-Synonymous mutations; $\pi_S$, nucleotide diversity by site for Synonymous mutations; $\pi_{NS}$, nucleotide diversity by site for Non-Synonymous mutations; $\pi(NS/S)$, Ratio of $\pi_{NS}$ to $\pi_{Syn}$.

**Table S4. Genetic diversity in the non-coding regions of the mitochondrial genome (842 bps)**

| | | *N* | *MD* | *H* | $H_d$ (%) | *S* | *Shared P.* | *Singl.* | $\pi$ (%) | $\vartheta_W$ (%) |
|---|---|---|---|---|---|---|---|---|---|---|
| Species | All | 63 | 13 | 41 | 96.8 | 114 | 107 | 7 | 4.62 | 3.16 |
| | FP | 12 | 8 | 7 | 83.3 | 10 | 5 | 5 | 0.36 | 0.40 |
| | BP | 3 | 5 | 3 | 100.0 | 5 | 0 | 5 | 0.40 | 0.40 |
| | V | 12 | 4 | 4 | 45.5 | 4 | 0 | 4 | 0.08 | 0.13 |
| | SP | 3 | 5 | 3 | 100.0 | 8 | 0 | 8 | 0.68 | 0.64 |
| | DP | 6 | 8 | 5 | 93.3 | 15 | 5 | 10 | 0.74 | 0.80 |
| | HP | 27 | 10 | 19 | 95.7 | 39 | 30 | 9 | 1.21 | 1.15 |
| HP | NAT | 4 | 10 | 4 | 100.0 | 5 | 1 | 4 | 0.32 | 0.33 |
| | IB | 3 | 8 | 2 | 66.7 | 1 | 0 | 1 | 0.08 | 0.08 |
| | MA | 3 | 8 | 3 | 100.0 | 5 | 0 | 5 | 0.40 | 0.40 |
| | BS | 3 | 8 | 3 | 100.0 | 2 | 0 | 2 | 0.16 | 0.16 |
| | NP | 14 | 7 | 7 | 84.6 | 11 | 8 | 3 | 0.45 | 0.42 |
| NP | NP1 | 10 | 7 | 4 | 71.1 | 6 | 1 | 5 | 0.19 | 0.26 |
| | NP2 | 4 | 6 | 2 | 66.7 | 2 | 1 | 1 | 0.14 | 0.13 |
| FP | YFP | 6 | 8 | 2 | 33.3 | 2 | 0 | 2 | 0.08 | 0.11 |
| | EAF | 5 | 8 | 4 | 90.0 | 6 | 1 | 5 | 0.31 | 0.35 |
| DP | DP2 | 5 | 8 | 4 | 90.0 | 7 | 4 | 3 | 0.43 | 0.41 |
| SP | SP2 | 2 | 5 | 2 | 100.0 | 4 | 0 | 4 | 0.35 | 0.35 |

N, Sample size; MD, Number of sites with missing data; *H*, number of haplotypes; $H_d$, haplotypic diversity; *S*, segregating sites; *Shared P.*, shared polymorphism; *Singl.*, singleton; $\pi$, average nucleotide diversity by site; $\theta_W$, theta from *S*. The meaning of the group acronyms is provided in Table S3.

**Table S5. Time (in million years) to the most recent common ancestor of lineages analyzed in this study.**

| Clade | Median age | 95% HPD |
|---|---|---|
| FP-V-BP-SP-DP-HP | 5.42 | 4.24-6.89 |
| V-BP-SP-DP-HP | 4.06 | 3.15-5.12 |
| V-BP-SP | 2.39 | 1.74-3.19 |
| BP-SP | 2.14 | 1.51-2.86 |
| DP-HP | 3.12 | 2.31-3.98 |
| FP | 0.49 | 0.31-0.78 |
| YFP-EAF | 0.22 | 0.14-0.35 |
| YFP | 0.06 | 0.03-0.11 |
| EAF | 0.13 | 0.08-0.21 |
| V | 0.04 | 0.02-0.07 |
| BP | 0.08 | 0.04-0.14 |
| SP | 0.43 | 0.27-0.67 |
| SP2 | 0.13 | 0.06-0.21 |
| DP | 0.48 | 0.32-0.77 |
| DP2 | 0.24 | 0.16-0.39 |
| HP | 0.86 | 0.76-0.90 |
| NP | 0.21 | 0.14-0.33 |
| NP1 | 0.1 | 0.06-0.15 |
| NP2 | 0.02 | 0.01-0.05 |
| NA | 0.41 | 0.31-0.55 |
| NAT | 0.22 | 0.14-0.31 |
| IB | 0.09 | 0.05-0.15 |
| MA | 0.06 | 0.03-0.11 |
| BS | 0.05 | 0.02-0.09 |

HPD = highest posterior density of the age of the lineage.
The meaning of the group acronyms is provided in Table S3.

### Supplementary Figures

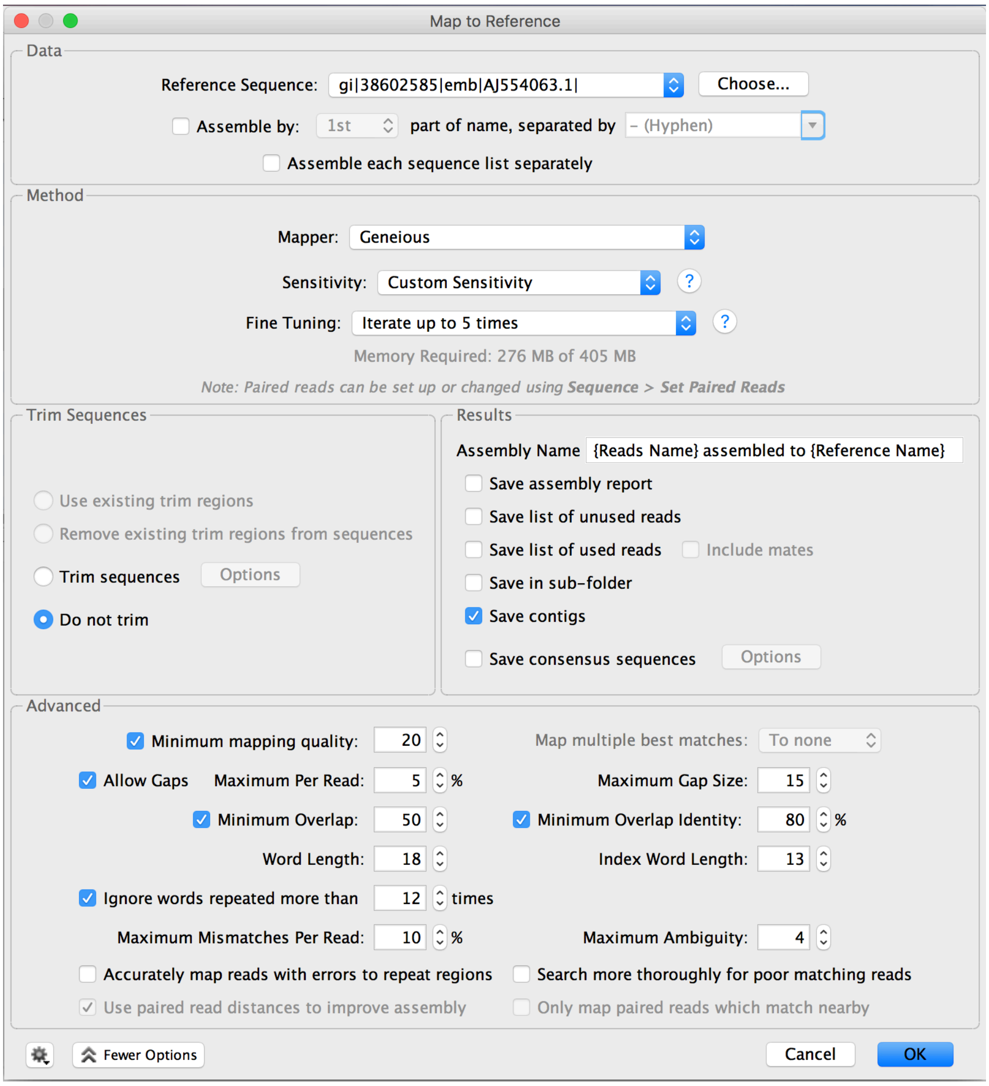

**Fig. S1. Geneious assembly parameters.**

Set of options used in Geneious v.8.1.9 to assemble the cleaned reads, including a first step where the reads are mapped against the harbor porpoise mitogenome reference (Arnason et al., 2004) and a second step to infer the consensus sequence from the mapped reads.

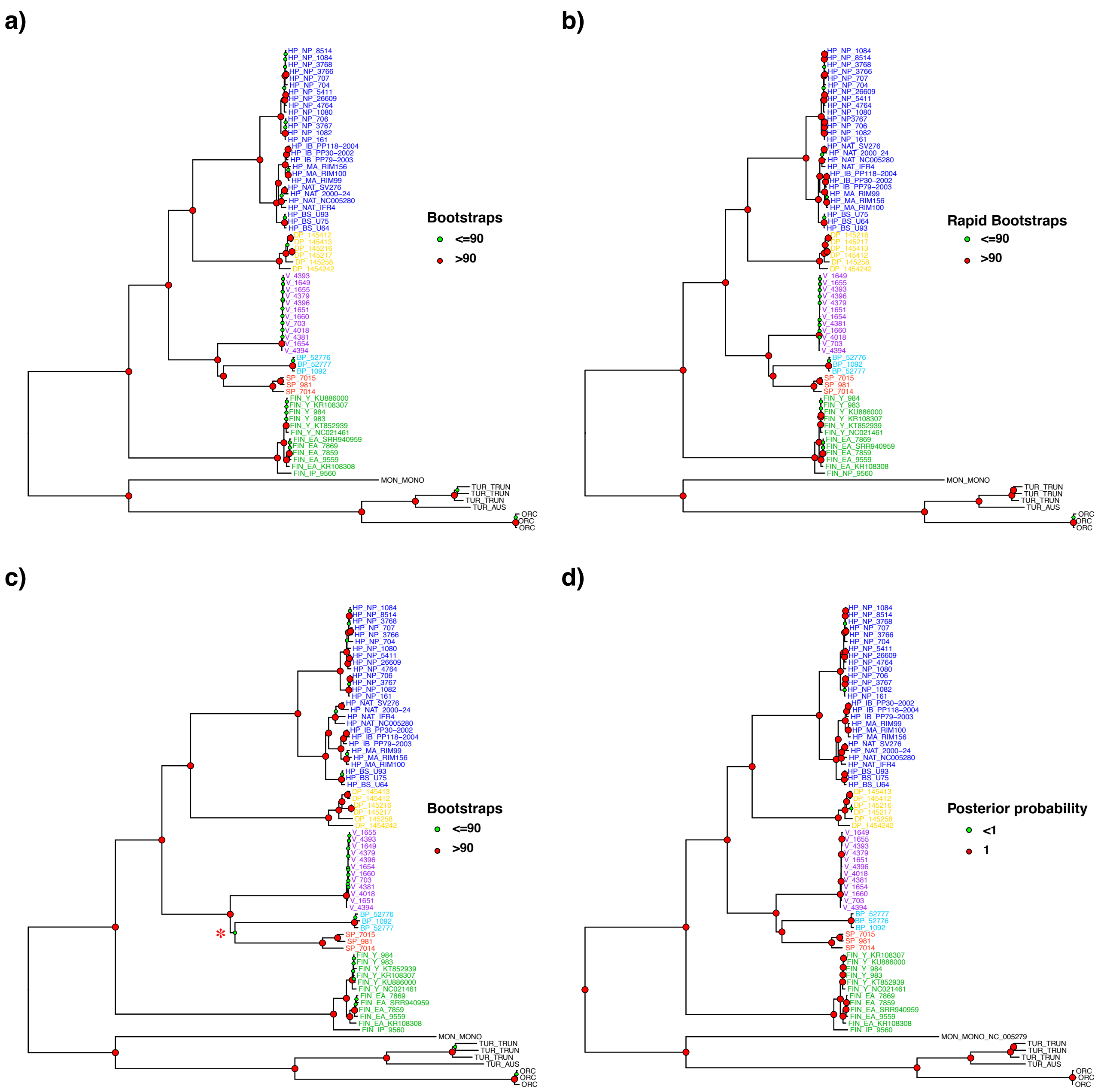

**Fig. S2. Mitochondrial phylogeny estimated using four different approaches.**

(a) Maximum likelihood phylogeny without partition and (b) with partitionning; (c) distance based (NJ) mitochondrial phylogeny; and (d) Bayesian mitochondrial phylogeny. The statistical support of each node for each method is indicated by the node color coding with the bootstrap support or posterior probability. The external branches and tip labels are colored according to the species. The tree is rooted with eight sequences from four closely related *Odontoceti* species (one *M. Monoceros,* three *T. truncatus,* one *T. australis* and three *O. orca* in black). The meaning of the group acronyms is provided in Table S3.

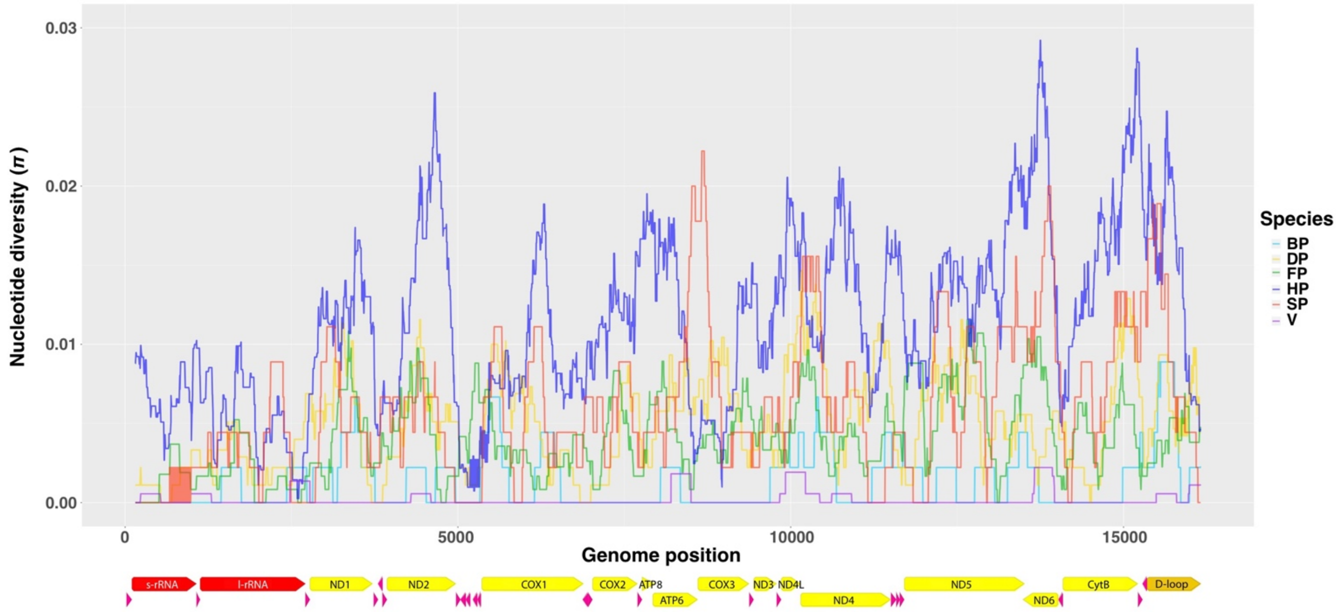

**Fig. S3. Nucleotide diversity (π) along the mitogenome of six species of porpoises.** The protein coding genes (yellow), rRNA (red), tRNA (pink) and D-loop (gold) are shown below the x-axis. Arrows indicate the direction of transcription. The meaning of the group acronyms is provided in Table S3.

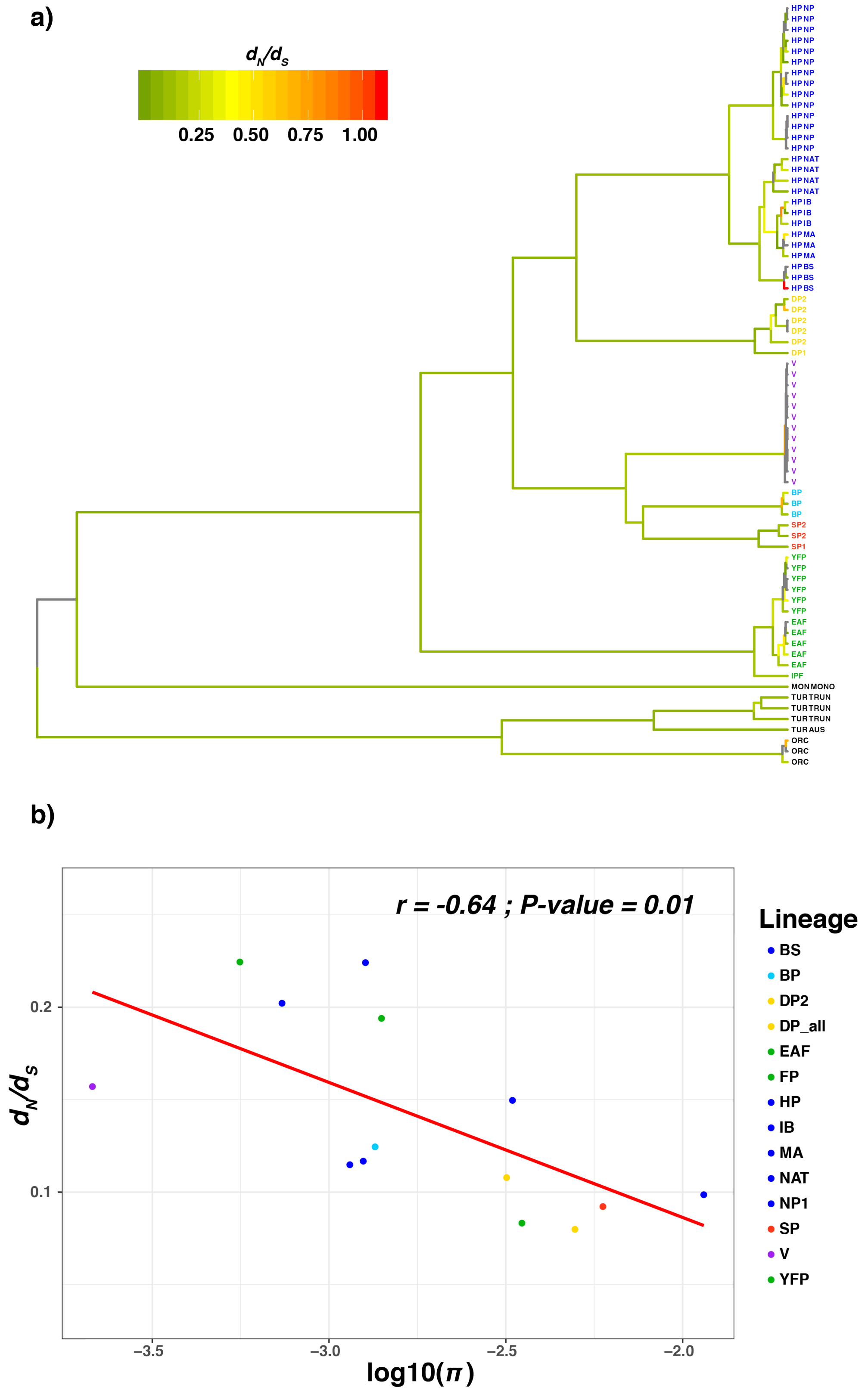

**Fig. S4. Evolution of the $d_N/d_S$ ratio in the porpoise family.**

(a) Maximum likelihood mitochondrial phylogeny in which the branches have been colored according to the $d_N/d_S$ ratio. (b) Relationship between mtDNA $d_N/d_S$ ratio and log scaled nucleotide diversity ($\pi$) across the porpoise family. The regression line is shown in red. The meaning of the group acronyms is provided in Table S3.

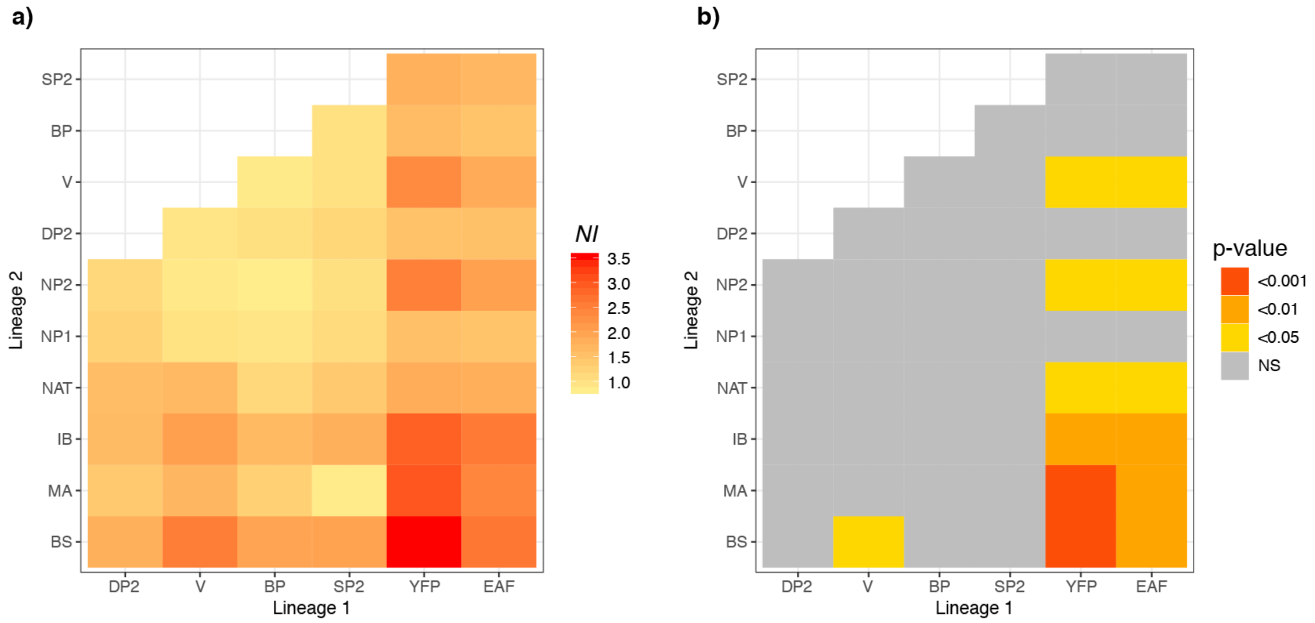

**Fig. S5. Heat map of neutrality index of McDonald-Kreitman (MK) tests between all pairwise interspecific lineages.**

(a) Pairwise Neutral Index (NI) and (b) *p*-values associated with the MK tests. The meaning of the group acronyms is provided in Table S3.
